# SAR11 Genome Atlas: a genome and gene catalog for functional profiling of the most abundant bacterial clade in the ocean

**DOI:** 10.64898/2026.08.15.744057

**Authors:** Satoshi Nishino, Kento Tominaga, Hajime Itoh, Kimiho Omae, Koji Hamasaki, Susumu Yoshizawa, Yuki Nishimura

## Abstract

The SAR11 clade, also known as the order *Pelagibacterales*, is among the most abundant bacterial lineages in the ocean and plays central roles in marine biogeochemical cycles. However, many SAR11 genes remain functionally uncharacterized, highlighting the need for a comprehensive, integrated catalog that supports genomic, functional, and ecological analyses across the clade. Here, we present the SAR11 Genome Atlas, an interactive ortholog group (OG)-centered web resource that integrates 542 SAR11 genomes, including all 132 cultured strain genomes, with functional annotations, synteny, phylogenetic distribution, metatranscriptomic expression, and predicted protein structure information. To demonstrate its utility, we used environmental expression profiles to identify OGs associated with high-latitude environments, recovering OGs known to be involved in cold adaptation and proposing a functional hypothesis for an uncharacterized protein. We further analyzed phylogenetic distribution patterns to identify mutually exclusive functional modules, including candidate alternative systems for Mn/Zn homeostasis and phosphate acquisition, and to examine their association with distinct oceanographic environments. Together, these case studies highlight the value of the SAR11 Genome Atlas for generating ecological hypotheses by providing environmental and phylogenetic context for genes of interest. Through a user-friendly web interface, the SAR11 Genome Atlas enables researchers to explore genomic, environmental, and structural information without specialized computational expertise. All data and analysis outputs are freely accessible online at [https://stsnsn.github.io/SAR11_Atlas/].

## Introduction

The SAR11 clade, also known as the order *Pelagibacterales*, plays central roles in global biogeochemical cycles owing to its remarkable abundance and widespread distribution in the ocean. This clade accounts for approximately 25% of prokaryotic cells in the surface ocean ^1^ and occurs in diverse aquatic environments, from polar to tropical waters and from surface waters to the deep ocean, as well as in estuarine and freshwater habitats ^2^. SAR11 bacteria are major consumers of dissolved organic matter and low-concentration substrates, contributing to carbon, nitrogen, sulfur, and phosphorus cycling ^3,4,2,5^. They also contribute to climate-relevant biogeochemical cycles through respiration and the transformation of dissolved organic carbon into climate-active gases, including methane and dimethyl sulfide ^2,6,7^. Therefore, elucidating the physiological characteristics and metabolic capabilities of SAR11 bacteria is essential for understanding their roles in marine ecosystems and global biogeochemical cycles.

The SAR11 clade possesses highly streamlined genomes (1.1 - 1.5 Mbp) ^8–10^. Nevertheless, the function of a substantial fraction of SAR11 genes remains uncharacterized because these genes lack detectable sequence similarity to experimentally characterized genes ^11^. Given the selective pressure associated with genome streamlining, many of these uncharacterized genes are likely to perform important functions in SAR11 physiology and ecological adaptation. Thus, in parallel with experimental studies that have revealed specialized molecular functions in SAR11 ^12,13,5,14^, bioinformatic analyses are increasingly important for inferring the potential functions of uncharacterized genes and prioritizing candidates for further experimental investigation. Such large-scale inference of gene function requires genomic resources that capture the diversity of the SAR11 clade and are readily accessible to a broad range of researchers regardless of their computational expertise.

Because sequence similarity alone is often insufficient for assigning functions to uncharacterized genes, combining complementary information from genomic context, phylogenetic distribution, environmental occurrence and expression, and protein structure can improve functional inference ^15,16,11^. A SAR11-specific resource should therefore integrate these diverse sources of information across the phylogenetic breadth of the clade. Unlike existing organism-specific databases such as EcoCyc and Cyanorak ^17,18^, a resource for SAR11 must accommodate extensive uncultured diversity and place greater emphasis on integrating diverse lines of evidence for functional inference rather than on curating existing functional knowledge. Although extensive cultivation efforts have yielded 132 isolate genomes ^19,8,20,21^, the currently recognized diversity of the SAR11 clade is far greater, as reflected by the 1,303 representative genomes registered in GTDB Release 11-RS232 ^22^. Moreover, SAR11 genomes exhibit extensive genetic diversity even among closely related populations. Such sequence heterogeneity complicates the reconstruction of metagenome-assembled genomes (MAGs), limiting the recovery of high-quality SAR11 MAGs from metagenomic data ^23,24^. Therefore, a reliable and comprehensive SAR11-specific database should integrate high-quality genomes from isolates and single-amplified genomes (SAGs) while covering the phylogenetic diversity of the clade ^25,26,8^. Existing resources provide several types of information relevant to functional inference, but these are distributed across separate resources and need to be integrated within a unified clade-wide framework. For example, marine metagenomic resources such as the Ocean Gene Atlas enable exploration of environmental sequence distributions ^27^, but are not genome-resolved databases. In contrast, STRING provides multiple types of protein associations that can be used for functional inference ^28^, but is designed neither for a specific lineage nor to integrate ecological information. Thus, a dedicated framework integrating genome-resolved, functional, and ecological information across the SAR11 clade is still needed.

Here, we present the SAR11 Genome Atlas [https://stsnsn.github.io/SAR11_Atlas/], a web-based interactive genome and gene resource for the SAR11 clade. The database integrates 542 SAR11 genomes, including all 132 publicly available isolate genomes together with SAGs and a limited number of MAGs. Genes are organized into ortholog groups (OGs) and linked to multiple layers of genomic and functional information, including synteny conservation, metatranscriptomic expression profiles, and predicted protein structures. By unifying genomic diversity within a single platform, the SAR11 Genome Atlas enables comprehensive exploration of their genomic features and gene functions. The SAR11 Genome Atlas was designed as an open and reusable resource following the Findable, Accessible, Interoperable, and Reusable (FAIR) principles ^29^, with curated data, analysis outputs, and interactive visualizations available through a web interface and as downloadable files. We anticipate that this resource will support SAR11 research across computational and experimental disciplines.

## Materials and Methods

### Genome dataset construction

We collected SAR11 and related genomes from previous studies ^19,30,31,10,25,32–36,4,37–42,26,43,44,21,20,8^ and submitted them to phylogenetic placement using GTDB-Tk v2.1.0 ^45^. We excluded genomes assigned to the order HIMB59 clade, also known as AEGEAN-169 and previously referred to as SAR11 clade V, because it is clearly resolved outside *Pelagibacterales* in current genome-based classifications ^46,47,8^. In contrast, clade IV genomes were retained based on their close phylogenetic relationship to SAR11 and their placement within the *Pelagibacterales* in the GTDB taxonomy, while their affiliation with the core SAR11 radiation remains debated ^8,26^. The high diversity among SAR11 populations makes it challenging to reconstruct reliable MAGs ^23^, and reconstructed SAR11 MAGs tend to lack diverse genomic islands ^48,49^. For these reasons, MAGs were generally excluded from the dataset, except when they represented subclades poorly represented by isolate genomes or SAGs, including freshwater and deep sea lineages.

The collected genomes were filtered following a previous study ^8^, using thresholds of >85% completeness and <10% contamination as estimated by CheckM2 v1.0.2 **(Figure S1, Tables S1 and S2)** ^50^. All 132 isolate genomes were retained, whereas MAGs serving as nomenclatural types under the SeqCode were evaluated individually when their estimated completeness fell near the threshold. The rationale for these exceptions and the treatment of genomes from subclade Ic are described in the **Supplementary Methods**. The final dataset comprised 542 SAR11 genomes, including 132 isolate genomes, 399 single-amplified genomes (SAGs), and 11 MAGs **(Figures 1A and 1B; Tables S1 and S2)**. The average carbon, nitrogen and sulfur content of amino acid residue side chains (C-ARSC, N-ARSC and S-ARSC, respectively) were calculated using quickARSC v0.5.2 ^51^, and genome GC content was calculated for each genome.

**Figure 1.**
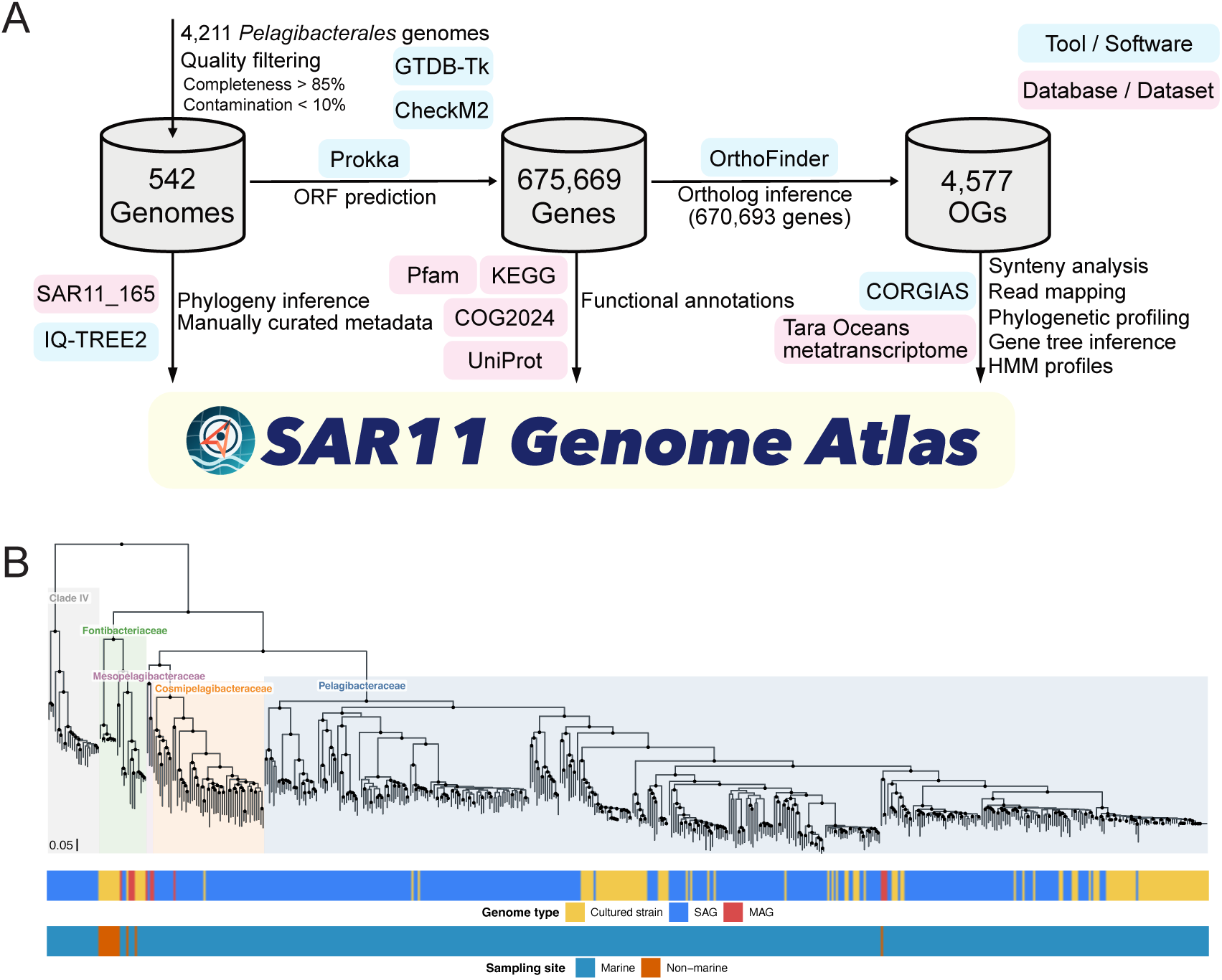
Construction and overview of the SAR11 Genome Atlas. (A) Workflow used to construct the SAR11 Genome Atlas. Following CheckM2 quality assessment and GTDB-Tk taxonomic screening, genes were predicted with Prokka and annotated using COG2024, KEGG Orthology, and Pfam. OrthoFinder assigned 670,693 of 675,669 proteins to 4,577 OGs. (B) Outgroup-rooted SAR11_165 phylogeny of the 542 SAR11 genomes inferred with IQ-TREE 2. The 20 alphaproteobacterial outgroup genomes used for rooting were removed before visualization. Shaded background colors indicate family-level SAR11 lineages. Black circles mark branches with ultrafast bootstrap support >= 95. The two tracks below the tree indicate genome type and sampling habitat (marine or non-marine). The scale bar represents 0.05 expected amino-acid substitutions per aligned site.

### Phylogeny inference

Heterogeneity in nucleotide composition and evolutionary rates across the *Alphaproteobacteria* genes can generate phylogenetic artifacts, such as long-branch attraction between *Pelagibacterales* and *Rickettsiales* ^52^. Because the phylogenetic position of *Pelagibacterales* within the *Alphaproteobacteria* remains unresolved, 20 isolate alphaproteobacterial genomes representing 10 orders were collected as outgroups. A concatenated alignment of 165 conserved SAR11 marker proteins (SAR11_165) was constructed by using hmmsearch from HMMER v3.4 (http://hmmer.org/), aligning the sequences with MAFFT in auto mode ^53^, and trimming the alignments with trimAl v1.5.rev1 using the -gt 0.50 option ^54^, following a previous study ^8^. The final alignment comprised 44,148 amino acid sites. Phylogenetic trees were inferred using IQ-TREE 2 v2.2.0.3 under the ModelFinder-selected VT+F+I+R10 model with 1,000 ultrafast bootstrap (UFBoot) and 1,000 Shimodaira-Hasegawa-like approximate likelihood ratio test (SH-aLRT) replicates ^55–57^. Phylogenetic trees were visualized using the R packages ggtree v3.10.1 and ggtreeExtra v1.12.0 ^58,59^.

Subclade, family, genus, and species assignments were harmonized using a hierarchical evidence framework. Direct taxonomic assignments from previously classified genomes and species representatives were prioritized, including the classification and taxonomic nomenclature of Freel et al. (2025) and the freshwater clade classifications of Lanclos et al. (2023) and Fernandes et al. (2025) ^8,20,35^. For genomes lacking direct assignments, species-level taxonomy was assigned when the ANI to a species representative genome was ≥95%, whereas family-, genus-, and subclade-level assignments were based on SAR11_165 marker-gene phylogenies. Detailed procedures for taxonomic assignment, confidence assessment, phylogenetic-depth calculation, and monophyly evaluation are provided in **Supplementary Text**.

### Ortholog group inference and functional annotation

All 675,669 protein coding sequences were predicted using Prokka v1.14.6 ^60^. Ortholog groups (OGs) were inferred using OrthoFinder v3.1.5 ^61^ with default parameters, resulting in 4,577 OGs containing 670,693 genes (99.3% of all predicted protein coding sequences). Predicted proteins were annotated against COG2024 using COGclassifier v2.0.0 (https://github.com/moshi4/COGclassifier), KEGG Orthology using KofamScan ^62^, and Pfam using PfamScan (https://github.com/aziele/pfam_scan), all of which were run with default settings. KofamScan applies profile-specific score thresholds designed to prioritize high confidence matches, which may result in more conservative KO assignments and lower annotation rates than those obtained using less stringent criteria. Functional annotations assigned to each OG were determined by majority vote, with the most frequently assigned annotation among member genes adopted as the representative annotation for the OG **(Table S3)**.

### Protein structure reference assignment

To associate SAR11 proteins with predicted structural information, all 675,669 predicted protein sequences were searched against UniProtKB release 2026_01 ^63^ using DIAMOND v2.1.10.164 ^64^. Swiss-Prot and TrEMBL were searched separately. Two searches were performed to identify close sequence matches (--id 85 --max-target-seqs 1) and more distant sequence homologs (--id 30 --max-target-seqs 10). UniProt accessions identified in either search were subsequently queried against the AlphaFold Protein Structure Database (AlphaFoldDB) ^65^, and matches with available structural models were classified into three categories: exact sequence matches, defined as 100% sequence identity with 100% query and subject coverage; close sequence matches, defined as at least 85% sequence identity with at least 80% query and subject coverage, excluding exact matches; and distant homologous matches, defined as at least 30% sequence identity, at least 80% query and subject coverage, and an E-value of at most 1 × 10⁻⁵. Exact and close sequence references were prioritized, and distant homologous references were used only for OGs without a qualifying exact or close sequence reference.

For the compact web table, candidate structure references were ranked by sequence identity, query coverage, subject coverage, bit score, and E-value; up to five high-ranking references were retained per OG. The complete DIAMOND search results are available through the web interface. The Protein Structure Viewer also provides links to the Foldseek web server for structural similarity searches ^66^.

### Genome synteny analysis

Synteny analysis was performed using GFF files generated by the Prokka pipeline ^60^. Following a previous study ^11^, neighboring genes were defined as the five genes located upstream and downstream on the same contig. For the web interface, all genes within this window of five upstream and five downstream genes were retained at the individual gene-occurrence level and used to visualize local synteny and operon-like gene organization across genomes. For network construction, gene pairs in which a given OG was found in the neighborhood of another OG in >= 75% of genomes containing that OG were considered neighboring gene pairs to minimize false positives arising from highly conserved SAR11 synteny ^10^. These filtered neighboring relationships were used to construct the neighboring gene network shown in the web interface.

### Phylogenetic profiling

To identify OGs with potentially related functions, we performed phylogenetic profiling based on their phylogenetic distributions across the SAR11 clade. Phylogenetic profiling compares the presence and absence patterns of genes across taxa, under the assumption that genes involved in related biological processes tend to exhibit correlated evolutionary histories ^67^. The analysis was performed using CORGIAS v1.1 ^68^ with an OG presence/absence matrix derived from OrthoFinder. Within the CORGIAS workflow, ancestral states were reconstructed using PastML based on the maximum-parsimony method ^69^. Significant OG associations were identified using the simultaneous evolutionary test implemented in CORGIAS ^68^, with positive and negative associations indicating concordant and opposing evolutionary patterns, respectively. *P*-values were adjusted using the Benjamini-Hochberg method, and associations with false discovery rate-adjusted *q-*values < 0.05 were retained. In the web interface, significant positive and negative associations obtained using the SAR11_165 phylogeny are displayed as red and blue edges, respectively.

### Tara Oceans metatranscriptome read mapping analysis

Sequence reads from the 509 metatranscriptomes obtained from the Tara Oceans samples ^70^ were mapped onto the 542 SAR11 genomes. We downloaded paired-end sequence data from NCBI SRA by SRA toolkit (https://github.com/ncbi/sra-tools), and then quality control was performed by using fastp with options -q 20 -n 10 -l 60 ^71^. The quality-controlled reads were mapped via bwa-mem2 ^72^. Transcripts per million (TPM) were calculated using a script from a previous study ^73^ and the summed TPM values of all genes belonging to each OG were used as the expression score.

### Literature metadata collection and citation network analysis

Literature information on SAR11 was obtained from PubMed and Web of Science on July 29, 2025, using the following search terms: “SAR11 OR Pelagibacter OR Pelagibacterales OR HTCC1062 OR Pelagiphage”. Citation count data for PubMed records were retrieved via DOI information using the Semantic Scholar API (Allen Institute for AI, https://www.semanticscholar.org). Literature networks were constructed using VOSviewer ^74^.

### Implementation of the SAR11 Genome Atlas web interface

The SAR11 Genome Atlas was implemented as a client-side web application using HTML, CSS, and JavaScript, with Bootstrap v4.3.1 for responsive page layout and interface styling (https://getbootstrap.com/). The interface includes modules for genome metadata browsing, OG annotation summaries, gene neighborhood and operon visualization, phylogenetic tree exploration, metatranscriptomic expression analysis, protein structure inspection, literature exploration, and dataset download. Tabular datasets were loaded in the browser from TSV/CSV files using Papa Parse (https://www.papaparse.com/) and rendered with DataTables (https://datatables.net/), enabling interactive searching, sorting, fixed headers, horizontal scrolling, and table export. Plotly.js was used for interactive charts and summary plots (https://plotly.com/javascript/), D3.js for SVG-based gene neighborhood visualization ^75^, and Cytoscape.js for neighboring gene and CORGIAS network visualization ^76^. Leaflet.js (https://leafletjs.com/) and Plotly Scattergeo (https://plotly.com) were used for geographic map visualization, including genome sampling locations and metatranscriptomic expression maps. Taxonium viewer was used for phylogenetic tree visualization ^77^, Mol* Viewer was used for interactive protein structure visualization ^78^, and VOSviewer v1.6.20 was used for literature-based network visualization ^74^. Additional custom JavaScript and CSS were used to implement theme switching, mobile sidebar navigation, search/autocomplete behavior, and synchronization between page-level controls and embedded viewers.

## Results and Discussion

### Construction of the SAR11 Genome Atlas

The SAR11 Genome Atlas was constructed as an OG-centered resource linking curated SAR11 genomes to functional annotation, genomic context, metatranscriptomic expression, phylogenetic distribution, and predicted protein structures **(Figure 1A)**. The current release integrates 542 genomes into 4,577 OGs comprising 670,693 protein-coding genes (99.3% of genes) **(Figure 1A)**. Functional annotations by sequence similarity search were available for 620,247 proteins (91.8%) through at least one of the COG2024, KEGG Orthology, or Pfam annotation systems. COG, KO, and Pfam annotations were detected for 593,170 (87.8%), 405,066 (60.0%), and 599,865 (88.8%) proteins, respectively. These annotations were integrated into downstream analyses across all 4,577 OGs, combining OG-level summaries, gene neighborhood and synteny relationships, phylogenetic profiling by CORGIAS, and Tara Oceans metatranscriptomic expression profiles.

Protein structure references from AlphaFoldDB were identified for 2,994 of the 4,577 OGs (65.4%), corresponding to 8,057 UniProt linked references ^63^. Exact sequence matches covered 1,409 OGs (30.8%), while additional close sequence matches covered 277 OGs (6.1%). Distant homologous matches extended coverage to a further 1,308 OGs (28.6%), leaving 1,583 OGs (34.6%) without a qualifying structural reference.

### Phylogenetic Placement of the family *Mesopelagibacteraceae*

The phylogenetic tree inferred from the SAR11_165 marker set recovered a topology generally consistent with the previous study ^8^ except for the placement of the deep sea lineage *Mesopelagibacteraceae*, which was recovered as sister to *Cosmipelagibacteraceae* with strong support **(UFBoot >= 95%, Figures 1B and S2)**. An additional bac120 tree, described in the **Supplementary Methods**, showed the same family-level topology, suggesting that the family-level classification in this study was robust to the choice of marker set. One likely explanation for the topological difference is the genome dataset used for phylogenetic inference. Freel et al. (2025) included five *Mesopelagibacteraceae* genomes, including the four SAGs reported by Thrash et al. (2014) ^25^, whereas our dataset contained only three genomes, including two MAGs, because the four SAGs, including the representative genome of *Mesopelagibacter profundi* (SAG, GCA_000504625.1), did not satisfy the genome-quality criteria assessed by CheckM2 **(Table S2)**. The evolutionary relationships among SAR11 lineages warrant further evaluation as additional high-quality genomes, particularly isolate genomes and high-quality SAGs from deep sea, become available (see **Scope, Limitations, and Future Directions**).

### Overview of the SAR11 Genome Atlas interface

The resulting datasets were released through the SAR11 Genome Atlas, an interactive web resource for exploring SAR11 gene conservation, genomic context, *in situ* gene expression, and predicted protein structure information (**Figure 2A-C**; https://stsnsn.github.io/SAR11_Atlas/). The interface was designed to connect complementary levels of biological information rather than present them as independent data tables. Representative views include the OG Information page, which summarizes OG annotation compositions and functional annotations **(Figure 2A)**; the Neighboring Genes page, which displays conserved local gene neighborhoods across genomes **(Figure 2B)**; and the Protein Structures page, which links SAR11 proteins to available AlphaFoldDB structure references **(Figure 2C)**.

**Figure 2.**
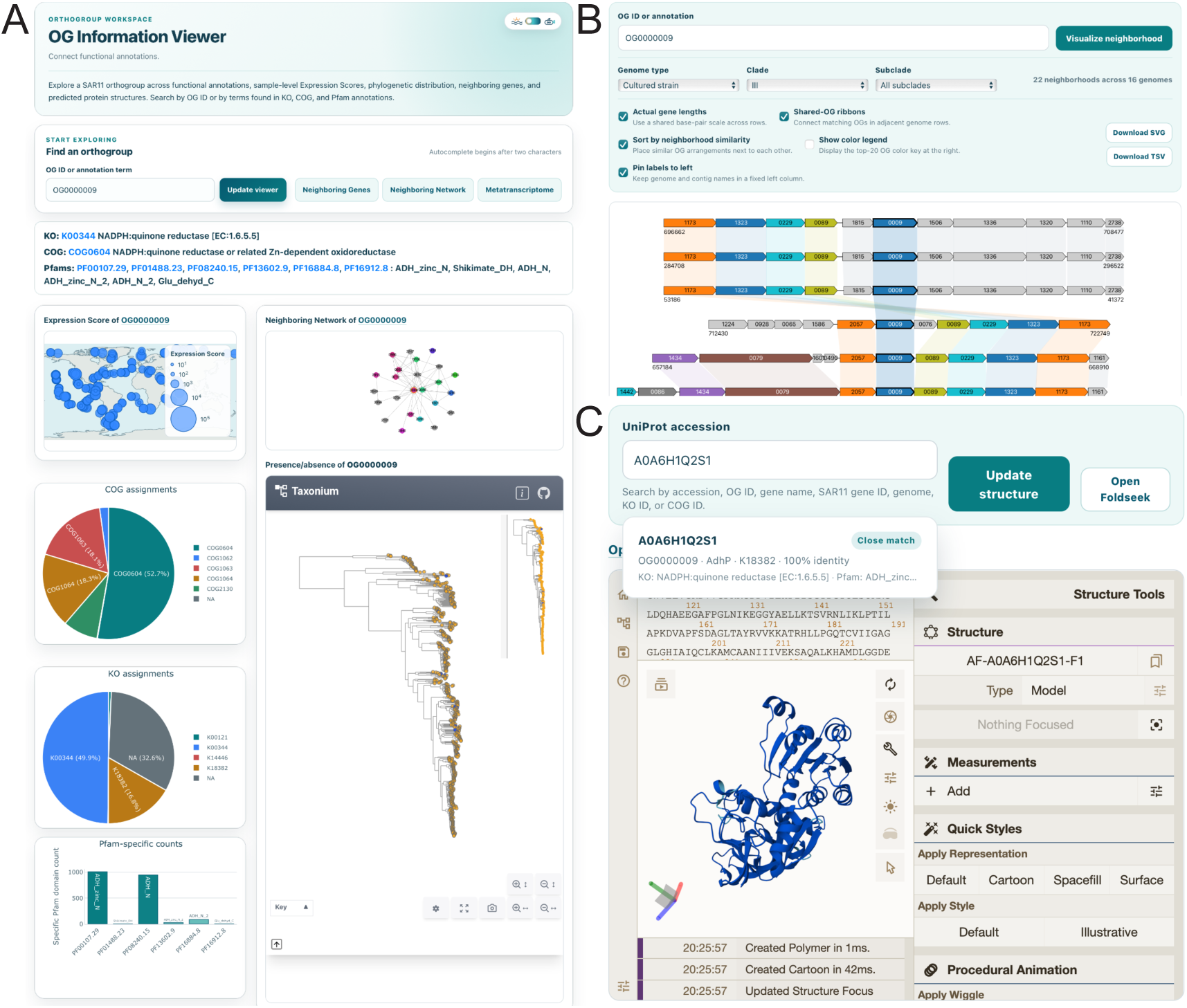
Examples of OG-centered exploration in the SAR11 Genome Atlas web interface. (A) OG Information page for OG0000009. The interface connects functional annotations with an OG-level Tara Oceans Expression Score map, COG, KEGG Orthology, and Pfam summaries, a neighboring-gene network, and the phylogenetic distribution of the OG in Taxonium. Links provide direct navigation to the neighboring-gene, network, metatranscriptome, and protein-structure views. (B) Neighboring Genes page for OG0000009. Gene-neighborhood diagrams can be filtered by genome type and taxonomic classification, sorted by neighborhood similarity, and displayed using shared gene-length and OG-color scales. Each row represents an occurrence of the query OG and its neighboring genes on the same contig; the underlying records can be downloaded as SVG or TSV files. (C) Protein Structures page for the UniProt accession A0A6H1Q2S1. Search suggestions connect UniProt accessions to OG, gene, genome, KO, COG, and Pfam annotations and distinguish close sequence matches from homologous structural references. An AlphaFoldDB model is displayed in the embedded Mol* viewer, and the selected structure can be submitted to Foldseek for structural similarity searches via direct link.

Users can begin with an OG or gene of interest and move across its functional annotations, phylogenetic distribution, neighboring gene context, co-occurrence or anti-occurrence relationships, metatranscriptomic expression profile, environmental correlations, and available structural references. Conversely, users can use genome metadata, sampling geography, genome type, or phylogenetic distribution as starting points to identify OGs associated with particular lineages or environmental contexts. This bidirectional organization supports both targeted investigation of known genes and discovery-oriented exploration of previously uncharacterized OGs. In addition to interactive exploration, the SAR11 Genome Atlas provides downloadable datasets for reproducible downstream analyses. Thus, the SAR11 Genome Atlas serves as both a visual exploration environment for hypothesis generation and an entry point to the underlying datasets.

Existing genomic and environmental resources provide complementary access to taxonomic, sequence-distribution, or organism-specific information. In contrast, the SAR11 Genome Atlas was designed specifically to connect genome-resolved OGs across cultured and uncultured SAR11 lineages with conserved gene neighborhoods, phylogenetically correlated OG associations, environmental expression profiles, and protein structure references. A comparison with representative existing resources is provided in **Table S4**. The comparison highlights that the principal contribution of the SAR11 Genome Atlas lies not only in providing individual data types but also in integrating them within a consistent OG-centered framework.

### Case study 1: High-latitude-associated expression pattern highlights potential cold adaptation in SAR11

To demonstrate the utility of the SAR11 Genome Atlas, we present an example application: identifying OGs whose transcriptional activity correlates with absolute latitude. Spearman rank correlations were calculated between the absolute latitude of each sampling site and the expression score **(Table S5)**. Using thresholds of Spearman’s ρ > 0.5, adjusted *q*-value < 0.05, and detectable expression in at least 10 samples, we identified 43 OGs with latitude-associated expression patterns **(Table S6)**. The strongest association was observed for OG0001080, encoding a CspA family cold shock protein **(ρ** = 0.743, Figure 3A).

**Figure 3.**
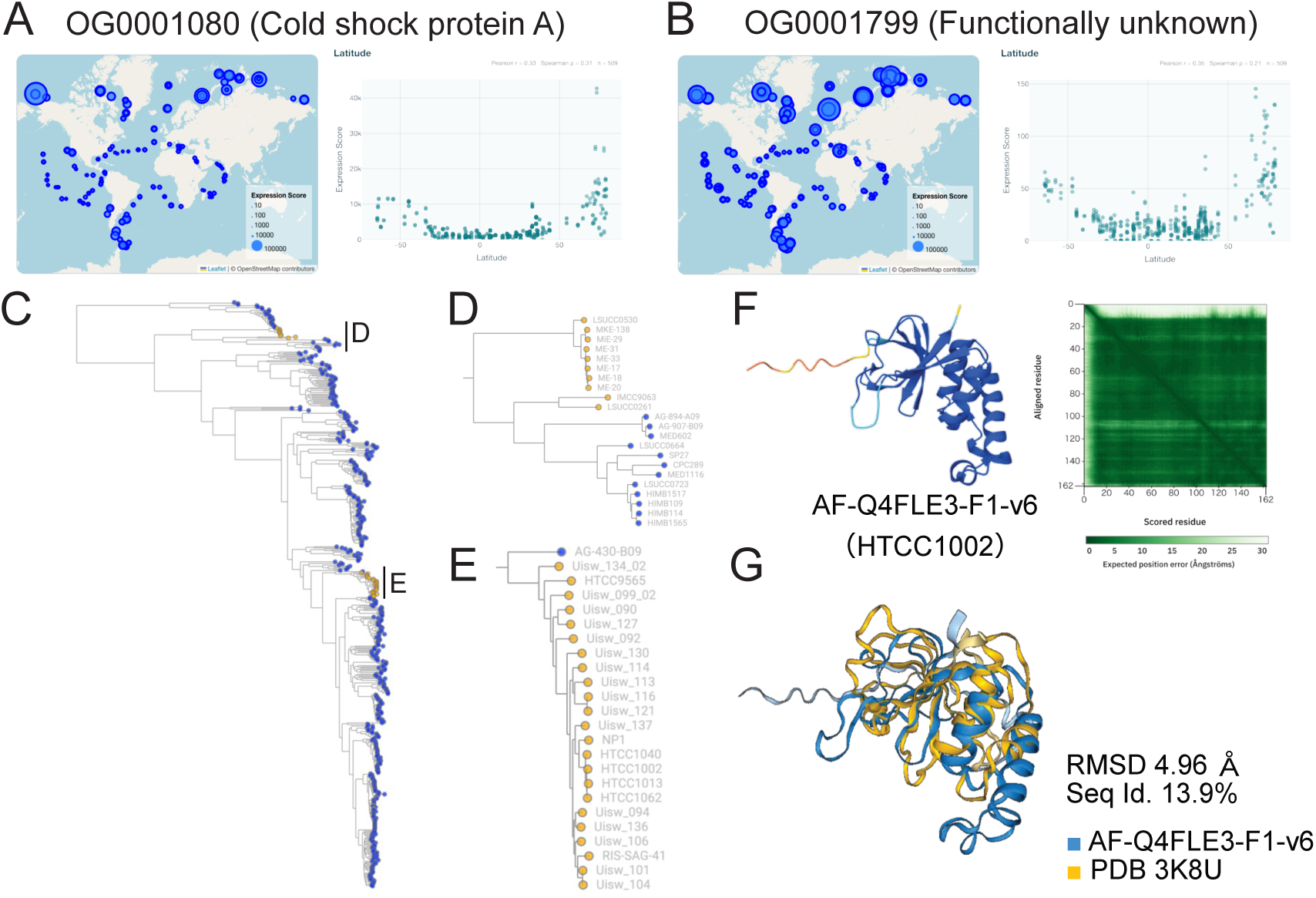
Case study 1: high-latitude expression patterns identify a known cold-response gene and support a functional hypothesis for OG0001799. (A) Geographic and latitudinal expression patterns of OG0001080, which encodes a CspA-family cold-shock protein. (B) Geographic and latitudinal expression patterns of the functionally uncharacterized OG0001799. In the maps, symbol size represents the OG-level Expression Score. Scatterplots show Expression Score against signed sampling latitude; the high-latitude screening analysis was performed using absolute latitude. (C) Phylogenetic distribution of OG0001799 across the 542-genome SAR11 tree. Tip-symbol colors indicate the presence (yellow) or absence (blue) of the OG. (D) Enlarged views of the freshwater subclade IIIb and the oligohaline-to-mesohaline subclade IIIa.3. (E) Enlarged view of the marine surface subclade Ia.1.I. (F) AlphaFoldDB model AF-Q4FLE3-F1-v6 as the representative protein of OG0001799 from HTCC1002, with the predicted aligned error matrix shown at right. (G) Structural superposition of AF-Q4FLE3-F1-v6 (blue) and the Foldseek-identified reference structure PDB 3K8U (yellow). The match had a Foldseek probability of 1.00, an amino-acid sequence identity of 13.9%, and a root mean square deviation of 4.96 angstroms, supporting structural similarity despite low sequence identity.

Polar oceans impose multiple physiological challenges on marine microbes, including low temperature and osmotic stress. For example, low temperatures reduce membrane fluidity, stabilize secondary structures of nucleic acids, and impair ribosome function and protein folding ^79,80^. Consistent with these established effects, several latitude-associated OGs have functions in RNA metabolism, ribosome biogenesis, compatible-solute transport, and fatty acid biosynthesis **(Figure S3)**. Together with the strong CspA signal, these results demonstrate that the SAR11 Genome Atlas recovers multiple independently interpretable components of bacterial cold adaptation from environmental expression data.

We next focused on functionally uncharacterized OGs among those associated with latitude and used the SAR11 Genome Atlas to investigate their potential functions. We selected OG0001799 as a representative example (**Figure 3B**). Its expression score increased with absolute latitude (ρ = 0.552), was negatively correlated with temperature (ρ = −0.564), and was positively correlated with oxygen concentration (ρ = 0.652). Collectively, OG0001799 showed preferential transcription under cold, oxygenated, high-latitude conditions. Its geographical expression pattern was concentrated primarily in surface ocean samples; however, its association with nominal sampling depth was weak and did not remain significant after multiple testing correction (ρ = −0.084, *q* = 0.072; **Table S5**).

The OG Information Viewer showed that OG0001799 was distributed across 37 genomes belonging to clades Ia and III **(Figure 3C–E)**. These included the marine surface clade Ia.1.I represented by *Pelagibacter ubiqueversans* HTCC1062, the freshwater clade IIIb represented by *Fontibacterium commune* LSUCC0530, and the oligohaline-to-mesohaline clade members of IIIa.3 represented by LSUCC0261. This distribution indicates that the OG is shared among the SAR11 clade occupying distinct aquatic environments. Structural inspection of a *Pelagibacter ubiqueversans* HTCC1002 homolog using the Protein Structure Viewer, followed by a Foldseek search, identified structural similarity to a peptidase domain (**Figure 3F-G**; Foldseek probability = 1.00, amino acid identity = 13.9%, and root mean square deviation = 4.96 Å). Because low temperature can impair protein folding and stability, the predicted peptidase-like structure suggests a possible role in protein quality control or turnover under cold conditions ^79,81^. This example demonstrates how the SAR11 Genome Atlas integrates environmental expression, phylogenetic distribution, genomic context, and structural similarity to generate testable functional hypotheses for previously uncharacterized genes.

### Case study 2: Phylogenetic profiling highlights contrasting functional strategies in the SAR11 clade

As a second case study, we demonstrate how the SAR11 Genome Atlas can be used to explore patterns of co-occurrence and mutual exclusivity among OGs as clues to functional relationships. Similar phylogenetic distributions may indicate shared pathways or functional processes, whereas mutually exclusive distributions may reflect functionally analogous systems or alternative lineage-specific strategies ^67,82–85^. We performed phylogenetic profiling to identify statistically supported OG associations in the SAR11 clade by using CORGIAS ^68^.

The resulting CORGIAS network contained 17,853 significant associations involving 2,982 OGs, including 17,631 positive and 222 negative associations. We used selected negative associations as case studies to demonstrate how the SAR11 Genome Atlas enables users to connect OG-level relationships with genomic, functional, ecological, and structural information. We examined Mn/Zn homeostasis and phosphate acquisition in detail as representative examples; the other modules involving iron uptake, oxidative stress defense, and heme biosynthesis are briefly described **(Table S7)**.

### Example 1: Alternative systems for Mn/Zn homeostasis

One of the clearest examples involved genes associated with transition metal homeostasis **(Figure 4A and E)**. OG0001606, annotated as a putative MntP-family Mn²⁺ efflux protein, showed significant anti-occurrence with three components of a Znu-like Mn²⁺/Zn²⁺ ABC uptake system: the substrate-binding protein ZnuA (OG0000963), permease ZnuB (OG0000964), and ATPase ZnuC (OG0000970; adjusted *q*-values = 8.20 × 10⁻^19^, 1.06 × 10⁻^20^, and 8.20 × 10⁻^19^, respectively). In contrast, the three ABC-transporter components strongly co-occurred with one another, consistent with their retention as a coherent uptake module. These distributions suggest that SAR11 genomes differentially retain an MntP-like efflux system or a Znu-like uptake system.

**Figure 4.**
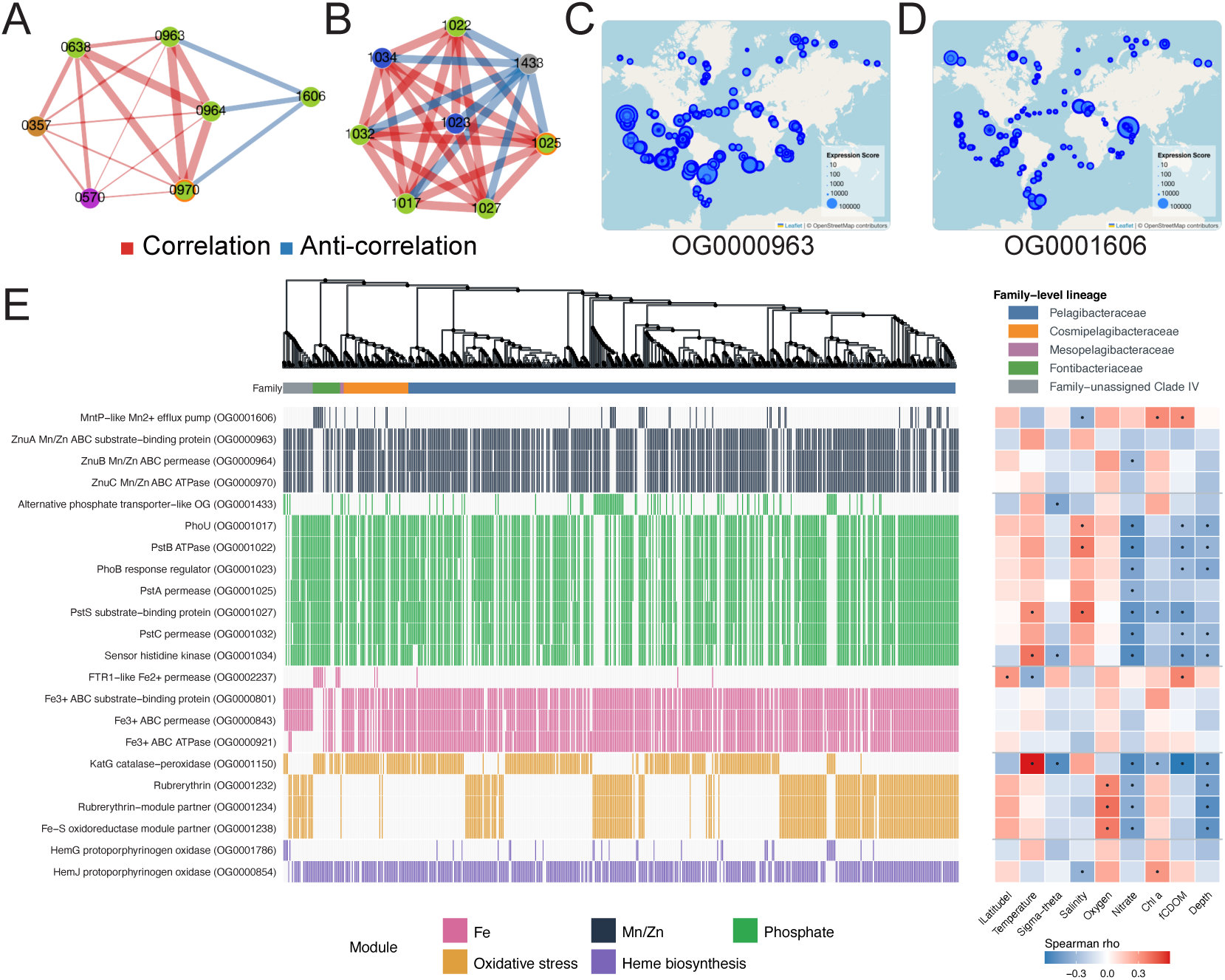
Case study 2: mutually exclusive SAR11 gene modules identified by CORGIAS. (A) CORGIAS subnetwork centered on the MntP-like Mn^2+^ efflux pump OG0001606 and components of the Znu-like Mn^2+^/Zn^2+^ ABC uptake system, including ZnuA (OG0000963), ZnuB (OG0000964), and ZnuC (OG0000970). (B) CORGIAS subnetwork connecting the alternative phosphate transporter-like OG0001433 with components of the canonical phosphate uptake and regulation system. In panels A and B, red edges indicate positive co-occurrence relationships, blue edges indicate anti-occurrence relationships, edge width reflects relationship strength, and node labels omit the "OG000" prefix for visibility. (C) Tara Oceans Expression Score map for the ZnuA substrate-binding protein OG0000963. (D) Tara Oceans Expression Score map for the MntP-like efflux pump OG0001606. Symbol size in panels C and D represents the OG-level Expression Score. (E) Joint phylogenetic and environmental summary of the 22 OGs selected for this case study. The upper tree shows the relationships among the 542 SAR11 genomes; black circles indicate branches with ultrafast bootstrap support >= 95, and the color track denotes family-level classification. Vertical marks show OG presence, with colors grouping OGs into Mn/Zn homeostasis, phosphate acquisition, iron acquisition, oxidative stress response, and heme biosynthesis modules. The heat map on the right shows Spearman rank correlations between each OG’s Expression Score and nine environmental variables across Tara Oceans metatranscriptomes. Black dots mark associations with absolute Spearman’s rho >= 0.3 and Benjamini-Hochberg-adjusted q < 0.05.

OG-level metatranscriptomic profiles suggested contrasting environmental conditions between the two systems **(Figure 4C–E and Table S8)**. The MntP-like OG0001606 was preferentially expressed in colder, lower-salinity, and more productive waters, with positive correlations with chlorophyll *a* and fCDOM. In contrast, the ZnuA-like substrate-binding component OG0000963 showed the broadly opposite pattern, with higher expression in warmer, more saline, and less productive waters. Because periplasmic substrate-binding proteins are frequently among the most abundant proteins expressed by SAR11 cells and underpin high-affinity acquisition of scarce substrates in oligotrophic waters ^5,14,86^, these opposing expression profiles support ecological differentiation between an MntP-like efflux strategy and a Znu-like uptake strategy. The permease and ATPase components showed generally compatible but weaker trends. The combined phylogenetic and expression patterns provide a testable hypothesis that different SAR11 lineages may use alternative strategies for Mn/Zn homeostasis.

### Example 2: A candidate alternative phosphate-acquisition system

A second prominent module involved genes associated with phosphate transport and regulation **(Figure 4B)**. OG0001433 showed significant mutual exclusion with seven components of a canonical Pst/Pho-associated system (*q*-values ranged from 8.72 × 10^-25^ to 8.07 × 10^-17^). These components strongly co-occurred with one another, supporting their interpretation as a coherent phosphate-uptake and regulatory module. Although OG0001433 was not assigned by the profile-based COGclassifier, KofamScan, or PfamScan workflows used here, the corresponding gene cluster was previously assigned to pitA (COG0306; phosphate/sulfate permease) using the sequence-based COG20/DIAMOND workflow implemented in anvi’o v8 ^87^. The same study further reported that genes of the high-affinity Pst/Pho system were core among genomes of coastal *Pelagibacteraceae* genera, whereas some offshore genomes lacking this system appeared to rely on *pitA* ^87^. Structural comparison of the representative OG0001433 protein using the Protein Structure Viewer and Foldseek revealed structural similarity to a sodium-dependent PiT family phosphate transporter (TmPiT; PDB 6L85; Foldseek probability = 1.00; amino acid identity = 16.8%; **Figure S4**) ^88^. The previous COG0306 assignment and structural similarity to TmPiT support a possible role in phosphate transport. Its mutual exclusion with the canonical Pst/Pho system further supports the hypothesis that OG0001433 is part of an alternative phosphate acquisition system. This case illustrates how genes that remain unannotated in our profile-based annotations can be linked to candidate functions through phylogenetic distribution and protein structure, consistent with independent sequence-based annotation.

The canonical Pst/Pho module was preferentially expressed in relatively warm, shallow, high-salinity, nitrate-depleted, and low-fCDOM samples **(Figures 4E and S4A; Table S8)**, consistent with activity in oligotrophic surface waters ^89,6,90,91^. OG0001433 showed weaker but partially overlapping associations, with relatively higher expression in warmer, shallower, lower salinity waters with higher chlorophyll *a*. This expression pattern is unexpected given the previous association of *pitA* with offshore genomes, which were linked to more oligotrophic conditions than coastal populations in the Hawaiian transect ^87^. However, the two analyses capture different ecological dimensions. The previous study examined lineage-specific gene content across Hawaiian coastal and offshore populations, whereas our analysis examined transcript distributions across globally distributed metatranscriptomic samples. The apparent difference may therefore reflect a distinction between the lineages in which the gene is retained and the environments in which its transcripts are relatively abundant. Because phosphate measurements were unavailable for the samples analyzed in this metatranscriptomic dataset, we could not determine whether OG0001433 expression was directly associated with phosphate availability. By enabling users to examine these genomic and expression patterns side by side within a single interface, the SAR11 Genome Atlas provides ecological context that is not apparent from genomic occurrence alone.

The contrasting genomic distributions and environmental expression profiles of the canonical Pst/Pho module and OG0001433 raise the possibility that they represent alternative phosphate acquisition systems. The Pst/Pho system is a well-established component of bacterial phosphate stress responses ^89,92^, whereas Pit/PiT family phosphate transporters are single component secondary transporters that couple transport to electrochemical ion gradients rather than ATP hydrolysis ^88,93^. Based on its previous COG0306 assignment and structural similarity to the Na^+^-dependent TmPiT, OG0001433 is a candidate Pit/PiT family phosphate transporter and may represent an energetically distinct alternative to the canonical Pst/Pho system. This example illustrates how the SAR11 Genome Atlas can integrate genomic distribution, environmental expression, and structural information to generate hypotheses about alternative phosphate acquisition strategies.

### Other examples of mutually exclusive functional modules

Three additional mutually exclusive modules further illustrated the diversity of candidate lineage-specific strategies: an FTR1-like Fe²⁺ permease versus an Fe³⁺ ABC uptake system, KatG versus a rubrerythrin-related oxidative stress defense module, and the non-homologous protoporphyrinogen oxidases HemG versus HemJ (Figure 4E; Table S8). Their distributions were associated with distinct phylogenetic and environmental patterns, suggesting alternative strategies for iron uptake, peroxide detoxification, and heme biosynthesis. These examples are summarized in **Table S7** and **Supplementary Text**. These additional modules further illustrate how the SAR11 Genome Atlas integrates gene distribution patterns with functional annotation and environmental information to reveal candidate lineage-specific metabolic strategies for experimental investigation.

### Scope, Limitations, and Future Directions

The SAR11 Genome Atlas provides an OG-centered framework that integrates phylogenetic, functional, environmental, and structural information across isolates and uncultured SAR11 lineages. By allowing users to move between environmental signals, phylogenetic distributions, genomic context, and protein structure, the Atlas supports both the functional investigation of uncharacterized genes and the ecological interpretation of known functions. Open access to curated datasets, analysis outputs, and reusable web resources provides a FAIR-oriented foundation for future comparative and ecological studies and for continued expansion as new SAR11 genomic and experimental data become available. Community feedback will also be important for correcting errors, incorporating new data, and maintaining links to external resources as the SAR11 Genome Atlas evolves.

The current version of the SAR11 Genome Atlas still has several limitations. Freshwater and deep sea SAR11 lineages remain underrepresented, and some environmental interpretations are constrained by incomplete metadata and indirect expression-based inference. Furthermore, phylogenetic profiling results may be influenced by the taxonomic scope and composition of the genome dataset, as gene distribution patterns depend on the lineages included in the analysis. Future updates could incorporate additional high-quality MAGs recovered through long-read metagenomics, additional deep sea SAGs ^94^, and expanded functional annotations supported by experimental studies ^12,5,95^. The phylogenetic placement and nomenclature of some SAR11 lineages may also require revision as broader *Alphaproteobacteria* phylogenies are refined, including the debated relationship between subclade IV and *Pelagibacterales*.

## Conclusions

Overall, the SAR11 Genome Atlas provides a foundation for connecting gene content to SAR11 ecology, physiology, and biogeochemistry across one of the most abundant bacterial groups in the ocean. By linking curated data products with interactive exploration, it offers a reusable framework for mining genes, identifying alternative metabolic strategies, and formulating experimentally testable hypotheses about SAR11 diversification across oceanographic gradients.

## Supporting information

Supplemental Figures S1 to S4 and legends for Supplemental Tables S1 to S8

## Supplementary Data

Supplementary Figures

Supplementary Text

## Acknowledgements

We would like to express our sincere gratitude to Naoki Konno for his suggestions and advice on the web interface development.

## Author contributions

**Satoshi Nishino** (Conceptualization [Lead], Methodology [Lead], Data curation [Lead], Formal analysis [Lead], Validation [equal], Visualization [Lead], Investigation [Lead], Funding acquisition [supporting], Writing—original draft [Lead]), **Kento Tominaga** (Formal analysis [supporting], Investigation [supporting], Supervision [equal], Writing—review & editing [equal]), **Hajime Itoh** (Validation [equal], Investigation [supporting], Data curation [supporting], Writing—review & editing [equal]), **Kimiho Omae** (Methodology [supporting], Writing—review & editing [equal]), **Koji Hamasaki** (Supervision [equal], Writing—review & editing [equal]), **Susumu Yoshizawa** (Supervision [equal], Writing—review & editing [equal]), **Yuki Nishimura** (Methodology [supporting], Resources [Lead], Validation [equal], Investigation [supporting], Funding acquisition [Lead], Supervision [equal], Writing—review & editing [equal])

## Data availability

The SAR11 Genome Atlas web interface is freely available at [https://stsnsn.github.io/SAR11_Atlas/] without login or registration. The source code, curated genome metadata, OG tables, functional annotations, phylogenetic trees, synteny relationships, phylogenetic profiling results, metatranscriptomic summaries, gene trees, HMM profiles, and protein structure reference tables are available from the Downloads page [https://stsnsn.github.io/SAR11_Atlas/html/download.html] and Zenodo [https://doi.org/10.5281/zenodo.21468730].

## References

1. Morris, R. M. et al. SAR11 clade dominates ocean surface bacterioplankton communities. Nature 420, 806–810 (2002).

2. Giovannoni, S. J. SAR11 Bacteria: The Most Abundant Plankton in the Oceans. Annu. Rev. Mar. Sci. 9, 231–255 (2017).

3. Malmstrom, R. R., Kiene, R. P., Cottrell, M. T. & Kirchman, D. L. Contribution of SAR11 Bacteria to Dissolved Dimethylsulfoniopropionate and Amino Acid Uptake in the North Atlantic Ocean. Appl. Environ. Microbiol. 70, 4129–4135 (2004).

4. Tsementzi, D. et al. SAR11 bacteria linked to ocean anoxia and nitrogen loss. Nature 536, 179–183 (2016).

5. Clifton, B. E., Alcolombri, U., Uechi, G.-I., Jackson, C. J. & Laurino, P. The ultra-high affinity transport proteins of ubiquitous marine bacteria. Nature 634, 721–728 (2024).

6. Carini, P., White, A. E., Campbell, E. O. & Giovannoni, S. J. Methane production by phosphate-starved SAR11 chemoheterotrophic marine bacteria. Nat. Commun. 5, 4346 (2014).

7. Sun, J. et al. The abundant marine bacterium Pelagibacter simultaneously catabolizes dimethylsulfoniopropionate to the gases dimethyl sulfide and methanethiol. Nat. Microbiol. 1, 16065 (2016).

8. Freel, K. C. et al. New SAR11 isolate genomes and global marine metagenomes resolve ecologically relevant units within the Pelagibacterales. Nat. Commun. 17, 328 (2025).

9. Giovannoni, S. J. et al. Genome Streamlining in a Cosmopolitan Oceanic Bacterium. Science 309, 1242–1245 (2005).

10. Grote, J. et al. Streamlining and Core Genome Conservation among Highly Divergent Members of the SAR11 Clade. mBio 3, 10.1128/mbio.00252-12 (2012).

11. Nishino, S. et al. Functional unknomics of the SAR11 clade reveal hidden genetic potential underlying adaptation to bottom-up and top-down pressures. mSystems e01026–26 (2026) doi:10.1128/msystems.01026-26.

12. Noell, S. E. & Giovannoni, S. J. SAR11 bacteria have a high affinity and multifunctional glycine betaine transporter. Environ. Microbiol. 21, 2559–2575 (2019).

13. Noell, S. E., Hellweger, F. L., Temperton, B. & Giovannoni, S. J. A Reduction of Transcriptional Regulation in Aquatic Oligotrophic Microorganisms Enhances Fitness in Nutrient-Poor Environments. Microbiol. Mol. Biol. Rev. 87, e00124–22 (2023).

14. Clifton, B. E., Akdavletov, B., Jain, P. & Laurino, P. Functional importance of a structurally encoded succinimide modification in a high-affinity solute-binding protein. 2026.06.18.732793 Preprint at 10.64898/2026.06.18.732793 (2026).

15. Pavlopoulos, G. A. et al. Unraveling the functional dark matter through global metagenomics. Nature 622, 594–602 (2023).

16. Baltoumas, F. A. et al. NMPFamsDB: a database of novel protein families from microbial metagenomes and metatranscriptomes. Nucleic Acids Res. gkad800 (2023) doi:10.1093/nar/gkad800.

17. Garczarek, L. et al. Cyanorak v2.1: a scalable information system dedicated to the visualization and expert curation of marine and brackish picocyanobacteria genomes. Nucleic Acids Res. 49, D667–D676 (2021).

18. Karp, P. D., Riley, M., Paley, S. M., Pellegrini-Toole, A. & Krummenacker, M. EcoCyc: Encyclopedia of Escherichia coli genes and metabolism. Nucleic Acids Res. 26, 50–53 (1998).

19. Rappé, M. S., Connon, S. A., Vergin, K. L. & Giovannoni, S. J. Cultivation of the ubiquitous SAR11 marine bacterioplankton clade. Nature 418, 630–633 (2002).

20. Fernandes, C. et al. Ecophysiology and global dispersal of the freshwater SAR11-IIIb genus Fontibacterium. Nat. Microbiol. 10, 2194–2206 (2025).

21. Sadler, M. C., Mino, S. & Morris, R. M. Complete genome sequences of 34 Arctic marine bacteria. Microbiol. Resour. Announc. 14, e00158–25 (2025).

22. Parks, D. H. et al. GTDB: an ongoing census of bacterial and archaeal diversity through a phylogenetically consistent, rank normalized and complete genome-based taxonomy. Nucleic Acids Res. 50, D785–D794 (2022).

23. Chang, T., Gavelis, G. S., Brown, J. M. & Stepanauskas, R. Genomic representativeness and chimerism in large collections of SAGs and MAGs of marine prokaryoplankton. Microbiome 12, 126 (2024).

24. Logares, R. Decoding populations in the ocean microbiome. Microbiome 12, 67 (2024).

25. Thrash, C. J. et al. Single-cell enabled comparative genomics of a deep ocean SAR11 bathytype. ISME J. 8, 1440–1451 (2014).

26. Haro-Moreno, J. M. et al. Ecogenomics of the SAR11 clade. Environ. Microbiol. 22, 1748–1763 (2020).

27. Vernette, C. et al. The Ocean Gene Atlas v2.0: online exploration of the biogeography and phylogeny of plankton genes. Nucleic Acids Res. 50, W516–W526 (2022).

28. Szklarczyk, D. et al. The STRING database in 2025: protein networks with directionality of regulation. Nucleic Acids Res. 53, D730–D737 (2025).

29. Wilkinson, M. D. et al. The FAIR Guiding Principles for scientific data management and stewardship. Sci. Data 3, 160018 (2016).

30. Stingl, U., Tripp, H. J. & Giovannoni, S. J. Improvements of high-throughput culturing yielded novel SAR11 strains and other abundant marine bacteria from the Oregon coast and the Bermuda Atlantic Time Series study site. ISME J. 1, 361–371 (2007).

31. Oh, H.-M. et al. Complete Genome Sequence of Strain IMCC9063, Belonging to SAR11 Subgroup 3, Isolated from the Arctic Ocean. J. Bacteriol. 193, 3379–3380 (2011).

32. Luo, H., Thompson, L. R., Stingl, U. & Hughes, A. L. Selection Maintains Low Genomic GC Content in Marine SAR11 Lineages. Mol. Biol. Evol. 32, 2738–2748 (2015).

33. Henson, M. W., Lanclos, V. C., Faircloth, B. C. & Thrash, J. C. Cultivation and genomics of the first freshwater SAR11 (LD12) isolate. ISME J. 12, 1846–1860 (2018).

34. Pachiadaki, M. G. et al. Charting the Complexity of the Marine Microbiome through Single-Cell Genomics. Cell 179, 1623–1635.e11 (2019).

35. Lanclos, V. C. et al. Ecophysiology and genomics of the brackish water adapted SAR11 subclade IIIa. ISME J. 17, 620–629 (2023).

36. Tully, B. J., Sachdeva, R., Graham, E. D. & Heidelberg, J. F. 290 metagenome-assembled genomes from the Mediterranean Sea: a resource for marine microbiology. PeerJ 5, e3558 (2017).

37. Hugerth, L. W. et al. Metagenome-assembled genomes uncover a global brackish microbiome. Genome Biol. 16, 279 (2015).

38. Cabello-Yeves, P. J. et al. Genomes of Novel Microbial Lineages Assembled from the Sub-Ice Waters of Lake Baikal. Appl. Environ. Microbiol. 84, e02132–17 (2017).

39. Jimenez-Infante, F. et al. Genomic characterization of two novel SAR11 isolates from the Red Sea, including the first strain of the SAR11 Ib clade. FEMS Microbiol. Ecol. 93, fix083 (2017).

40. Haro-Moreno, J. M. et al. Fine metagenomic profile of the Mediterranean stratified and mixed water columns revealed by assembly and recruitment. Microbiome 6, 128 (2018).

41. Zhao, Y. et al. Pelagiphages in the Podoviridae family integrate into host genomes. Environ. Microbiol. 21, 1989–2001 (2019).

42. Martinez-Hernandez, F. et al. Single-cell genomics uncover Pelagibacter as the putative host of the extremely abundant uncultured 37-F6 viral population in the ocean. ISME J. 13, 232–236 (2019).

43. Morris, R. M., Cain, K. R., Hvorecny, K. L. & Kollman, J. M. Lysogenic host– virus interactions in SAR11 marine bacteria. Nat. Microbiol. 5, 1011–1015 (2020).

44. Ruiz-Perez, C. A. et al. Description of Candidatus Mesopelagibacter carboxydoxydans and Candidatus Anoxipelagibacter denitrificans: Nitrate-reducing SAR11 genera that dominate mesopelagic and anoxic marine zones. Syst. Appl. Microbiol. 44, 126185 (2021).

45. Chaumeil, P.-A., Mussig, A. J., Hugenholtz, P. & Parks, D. H. GTDB-Tk v2: memory friendly classification with the genome taxonomy database. Bioinforma. Oxf. Engl. 38, 5315–5316 (2022).

46. Viklund, J., Martijn, J., Ettema, T. J. G. & Andersson, S. G. E. Comparative and Phylogenomic Evidence That the Alphaproteobacterium HIMB59 Is Not a Member of the Oceanic SAR11 Clade. PLOS ONE 8, e78858 (2013).

47. Getz, E. W. et al. The AEGEAN-169 clade of bacterioplankton is synonymous with SAR11 subclade V (HIMB59) and metabolically distinct. mSystems 8, e00179–23 (2023).

48. Rodriguez-Valera, F. et al. Explaining microbial population genomics through phage predation. Nat. Rev. Microbiol. 7, 828–836 (2009).

49. Molina-Pardines, C., Haro-Moreno, J. M., Rodriguez-Valera, F. & López-Pérez, M. Extensive paralogism in the environmental pangenome: a key factor in the ecological success of natural SAR11 populations. Microbiome 13, 41 (2025).

50. Chklovski, A., Parks, D. H., Woodcroft, B. J. & Tyson, G. W. CheckM2: a rapid, scalable and accurate tool for assessing microbial genome quality using machine learning. Nat. Methods 20, 1203–1212 (2023).

51. Nishino, S., Tominaga, K., Nishimura, Y. & Yoshizawa, S. quickARSC: standalone package and web interface for profiling elemental stoichiometry of proteomes. Microbiol. Resour. Announc. 15, e00031–26 (2026).

52. Muñoz-Gómez, S. A. et al. An updated phylogeny of the Alphaproteobacteria reveals that the parasitic Rickettsiales and Holosporales have independent origins. eLife 8, e42535 (2019).

53. Katoh, K. & Standley, D. M. MAFFT Multiple Sequence Alignment Software Version 7: Improvements in Performance and Usability. Mol. Biol. Evol. 30, 772–780 (2013).

54. Capella-Gutiérrez, S., Silla-Martínez, J. M. & Gabaldón, T. trimAl: a tool for automated alignment trimming in large-scale phylogenetic analyses. Bioinformatics 25, 1972–1973 (2009).

55. Kalyaanamoorthy, S., Minh, B. Q., Wong, T. K. F., von Haeseler, A. & Jermiin, L. S. ModelFinder: fast model selection for accurate phylogenetic estimates. Nat. Methods 14, 587–589 (2017).

56. Hoang, D. T., Chernomor, O., von Haeseler, A., Minh, B. Q. & Vinh, L. S. UFBoot2: Improving the Ultrafast Bootstrap Approximation. Mol. Biol. Evol. 35, 518– 522 (2018).

57. Minh, B. Q. et al. IQ-TREE 2: New Models and Efficient Methods for Phylogenetic Inference in the Genomic Era. Mol. Biol. Evol. 37, 1530–1534 (2020).

58. Xu, S. et al. ggtreeExtra: Compact Visualization of Richly Annotated Phylogenetic Data. Mol. Biol. Evol. 38, 4039–4042 (2021).

59. Yu, G., Smith, D. K., Zhu, H., Guan, Y. & Lam, T. T.-Y. ggtree: an R package for visualization and annotation of phylogenetic trees with their covariates and other associated data. Methods Ecol. Evol. 8, 28–36 (2017).

60. Seemann, T. Prokka: rapid prokaryotic genome annotation. Bioinformatics 30, 2068–2069 (2014).

61. Emms, D. M., Liu, Y., Belcher, L., Holmes, J. & Kelly, S. OrthoFinder: improved phylogenetic orthology inference with enhanced accuracy and scalability. Nat. Methods 23, 1327–1333 (2026).

62. Aramaki, T. et al. KofamKOALA: KEGG Ortholog assignment based on profile HMM and adaptive score threshold. Bioinformatics 36, 2251–2252 (2020).

63. The UniProt Consortium. UniProt: the Universal Protein Knowledgebase in 2025. Nucleic Acids Res. 53, D609–D617 (2025).

64. Buchfink, B., Reuter, K. & Drost, H.-G. Sensitive protein alignments at tree-of-life scale using DIAMOND. Nat. Methods 18, 366–368 (2021).

65. Varadi, M. et al. AlphaFold Protein Structure Database: massively expanding the structural coverage of protein-sequence space with high-accuracy models. Nucleic Acids Res. 50, D439–D444 (2022).

66. van Kempen, M. et al. Fast and accurate protein structure search with Foldseek. Nat. Biotechnol. 1–4 (2023) doi:10.1038/s41587-023-01773-0.

67. Pellegrini, M., Marcotte, E. M., Thompson, M. J., Eisenberg, D. & Yeates, T. O. Assigning protein functions by comparative genome analysis: Protein phylogenetic profiles. Proc. Natl. Acad. Sci. 96, 4285–4288 (1999).

68. Nishimura, Y., Omae, K., Tominaga, K. & Iwasaki, W. CORGIAS: identifying correlated gene pairs by considering evolutionary history in a large-scale prokaryotic genome dataset. NAR Genomics Bioinforma. 7, lqaf182 (2025).

69. Ishikawa, S. A., Zhukova, A., Iwasaki, W. & Gascuel, O. A Fast Likelihood Method to Reconstruct and Visualize Ancestral Scenarios. Mol. Biol. Evol. 36, 2069– 2085 (2019).

70. Salazar, G. et al. Gene Expression Changes and Community Turnover Differentially Shape the Global Ocean Metatranscriptome. Cell 179, 1068–1083.e21 (2019).

71. Chen, S., Zhou, Y., Chen, Y. & Gu, J. fastp: an ultra-fast all-in-one FASTQ preprocessor. Bioinformatics 34, i884–i890 (2018).

72. Vasimuddin, Md., Misra, S., Li, H. & Aluru, S. Efficient Architecture-Aware Acceleration of BWA-MEM for Multicore Systems. in 2019 IEEE International Parallel and Distributed Processing Symposium (IPDPS) 314–324 (2019). doi:10.1109/IPDPS.2019.00041.

73. Nishimura, Y. et al. Environmental Viral Genomes Shed New Light on Virus-Host Interactions in the Ocean. mSphere 2, 10.1128/msphere.00359-16 (2017).

74. van Eck, N. J. & Waltman, L. Software survey: VOSviewer, a computer program for bibliometric mapping. Scientometrics 84, 523–538 (2010).

75. Bostock, M., Ogievetsky, V. & Heer, J. D3 Data-Driven Documents. IEEE Trans. Vis. Comput. Graph. 17, 2301–2309 (2011).

76. Franz, M. et al. Cytoscape.js: a graph theory library for visualisation and analysis. Bioinformatics 32, 309–311 (2016).

77. Sanderson, T. Taxonium, a web-based tool for exploring large phylogenetic trees. eLife 11, e82392 (2022).

78. Sehnal, D. et al. Mol* Viewer: modern web app for 3D visualization and analysis of large biomolecular structures. Nucleic Acids Res. 49, W431–W437 (2021).

79. D’Amico, S. et al. Psychrophilic microorganisms: challenges for life. EMBO Rep. 7, 12 (2006).

80. Zhang, Y. & Gross, C. A. Cold Shock Response in Bacteria. Annu. Rev. Genet. 55, 377–400 (2021).

81. Merdanovic, M., Clausen, T., Kaiser, M., Huber, R. & Ehrmann, M. Protein Quality Control in the Bacterial Periplasm. Annu. Rev. Microbiol. 65, 149–168 (2011).

82. Morett, E. et al. Systematic discovery of analogous enzymes in thiamin biosynthesis. Nat. Biotechnol. 21, 790–795 (2003).

83. Kensche, P. R., van Noort, V., Dutilh, B. E. & Huynen, M. A. Practical and theoretical advances in predicting the function of a protein by its phylogenetic distribution. J. R. Soc. Interface 5, 151–170 (2007).

84. Kumagai, Y. et al. Solar-panel and parasol strategies shape the proteorhodopsin distribution pattern in marine Flavobacteriia. ISME J. 12, 1329–1343 (2018).

85. Tominaga, K. et al. Prevalence of Viral Frequency-Dependent Infection in Coastal Marine Prokaryotes Revealed Using Monthly Time Series Virome Analysis. mSystems 8, e00931–22 (2023).

86. Sowell, S. M. et al. Transport functions dominate the SAR11 metaproteome at low-nutrient extremes in the Sargasso Sea. ISME J. 3, 93–105 (2009).

87. Tucker, S. J., Freel, K. C., Eren, A. M. & Rappé, M. S. Habitat-specificity in SAR11 is associated with a few genes under high selection. ISME J. 19, wraf216 (2025).

88. Tsai, J.-Y. et al. Structure of the sodium-dependent phosphate transporter reveals insights into human solute carrier SLC20. Sci. Adv. 6, eabb4024 (2020).

89. Rao, N. N. & Torriani, A. Molecular aspects of phosphate transport in Escherichia coli. Mol. Microbiol. 4, 1083–1090 (1990).

90. Zubkov, M. V., Martin, A. P., Hartmann, M., Grob, C. & Scanlan, D. J. Dominant oceanic bacteria secure phosphate using a large extracellular buffer. Nat. Commun. 6, 7878 (2015).

91. Zhu, W.-J. et al. Structural and molecular basis for phosphate recognition by SAR11 bacteria. mBio 16, e01654–25 (2025).

92. Kim, S.-K. et al. Dual Transcriptional Regulation of the Escherichia coli Phosphate-Starvation-Inducible psiE Gene of the Phosphate Regulon by PhoB and the Cyclic AMP (cAMP)-cAMP Receptor Protein Complex. J. Bacteriol. 182, 5596–5599 (2000).

93. Harris, R. M., Webb, D. C., Howitt, S. M. & Cox, G. B. Characterization of PitA and PitB from Escherichia coli. J. Bacteriol. 183, 5008–5014 (2001).

94. Chang, T. et al. Single-cell-resolved genome atlas of prokaryoplankton inhabiting the ocean’s interior. Cell (2026).

95. Zhang, Y. et al. The functional proteome landscape of a cosmopolitan oceanic bacterium. Protein Cell pwag074 (2026) doi:10.1093/procel/pwag074.

