## Supplemental Figures S1 to S4 and legends for Supplemental Tables S1 to S8 for "SAR11 Genome Atlas: a genome and gene catalog for functional profiling of the most abundant bacterial clade in the ocean"

### Graphical Abstract

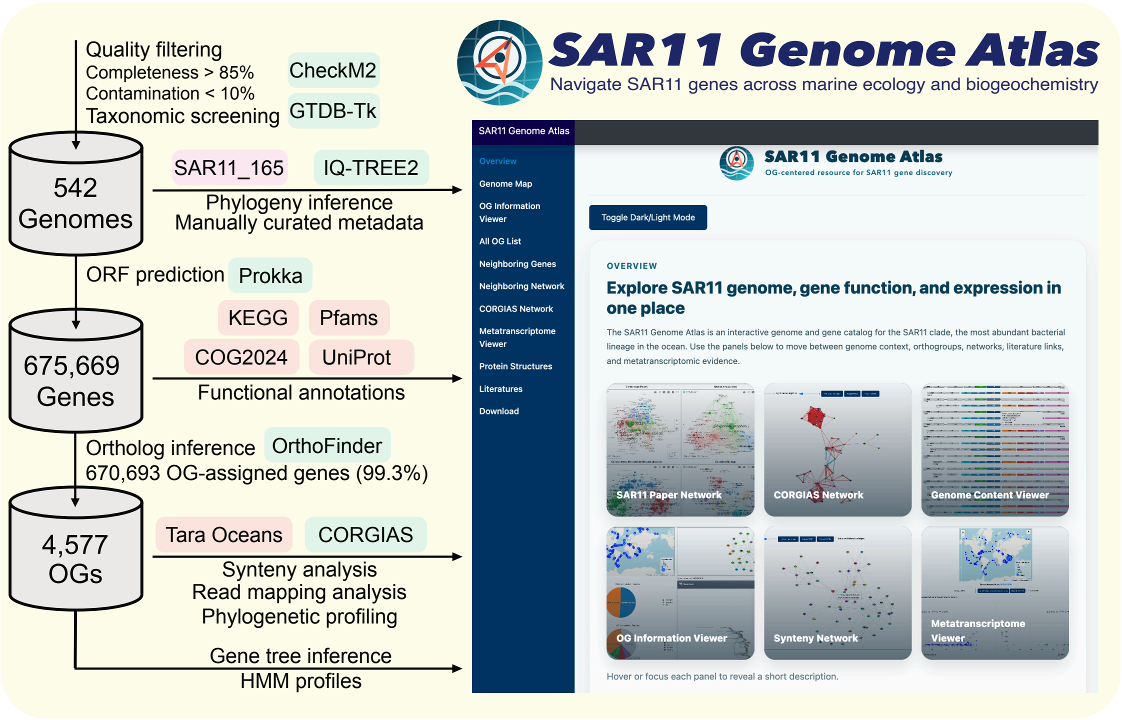

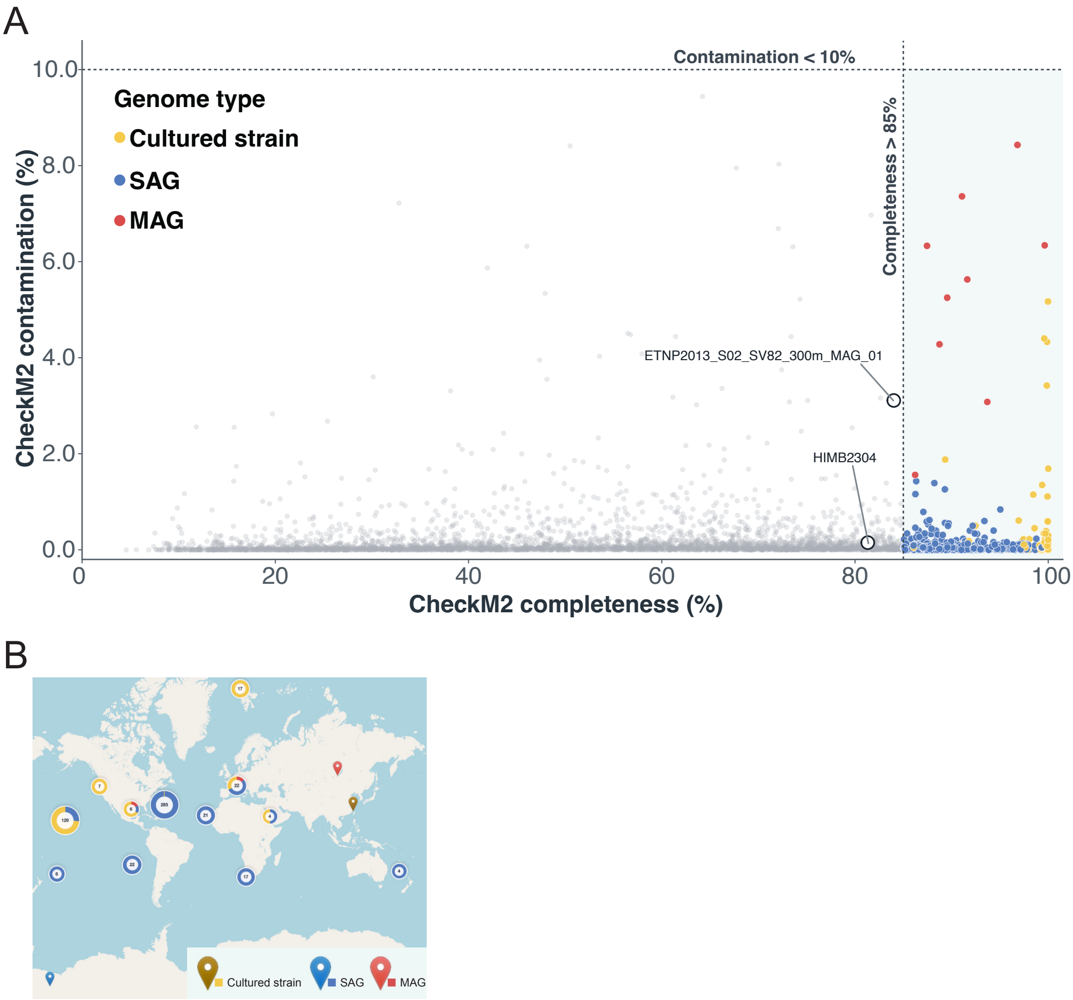


**Supplementary Figure S1.** (A) Genome quality and selection. CheckM2 completeness and contamination estimates are shown for 4,211 screened candidate genomes. The 542 genomes retained in the Atlas are colored by genome type (132 cultured strains, 399 single-amplified genomes, and 11 metagenome-assembled genomes); genomes not retained are shown in gray. Dashed lines indicate the principal selection thresholds of completeness >85% and contamination <10%. HIMB2304 and ETNP2013_S02_SV82_300m_MAG_01 are labeled as documented inclusion exceptions. (B) Geographic distribution of the screened SAR11 genomes. Symbols indicate cultured strains, SAGs, and MAGs, and clustered symbols show the number and composition of genomes sampled from nearby locations.

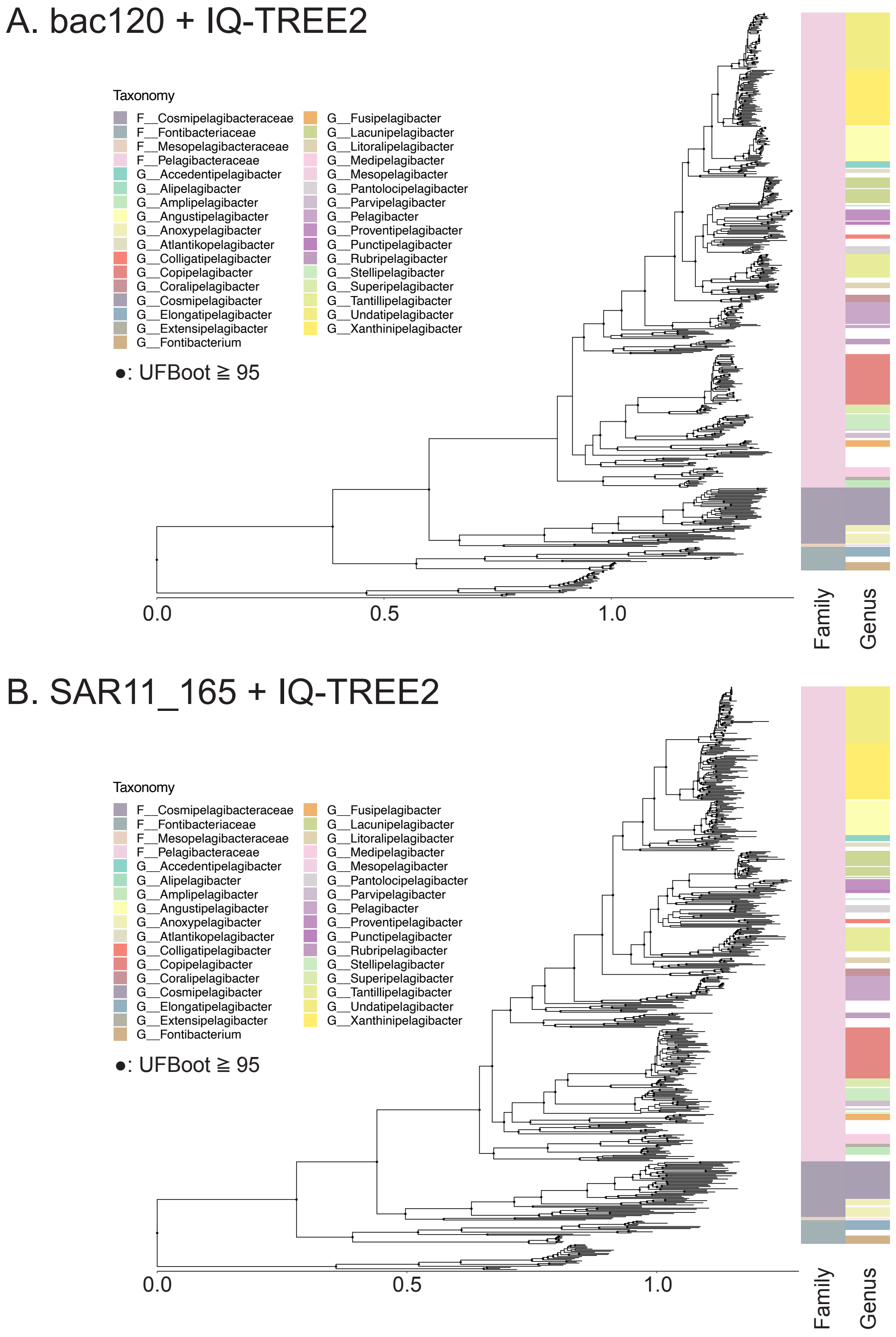


**Supplementary Figure S2. Phylogenies of the SAR11 clade used in this study.** (A) Phylogenetic tree based on the GTDB bacterial marker gene set (bac120). (B) Phylogenetic tree based on the SAR11-specific 165 single-copy core-gene set (SAR11_165). The adjacent heatmaps show family and genus assignments. Black points on the trees indicate internal nodes with ultrafast bootstrap support values of at least 95. Taxonomic names beginning with “F__” denote families, whereas those beginning with “G__” denote genera.

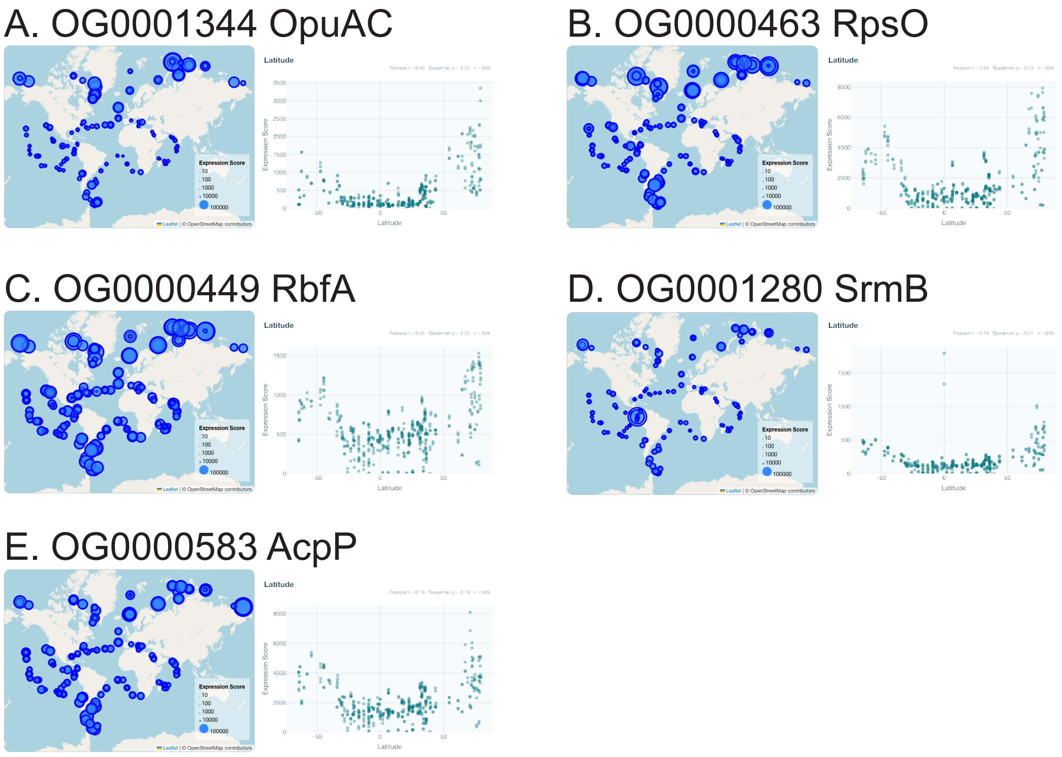


**Supplementary Figure S3. Additional high-latitude-associated OGs with functions related to osmoprotection, translation, RNA remodeling, and membrane-lipid biosynthesis.**

Geographic Expression Score maps and scatterplots of Expression Score against signed sampling latitude are shown for (A) OG0001344, encoding the compatible-solute-binding protein OpuAC; (B) OG0000463, encoding ribosomal protein RpsO; (C) OG0000449, encoding ribosome-binding factor A (RbfA); (D) OG0001280, encoding the DEAD-box RNA helicase SrmB; and (E) OG0000583, encoding acyl carrier protein AcpP. Symbol size on each map represents the OG-level Expression Score. These OGs were identified by screening correlations with absolute latitude; signed latitude is displayed to show expression patterns in both hemispheres.

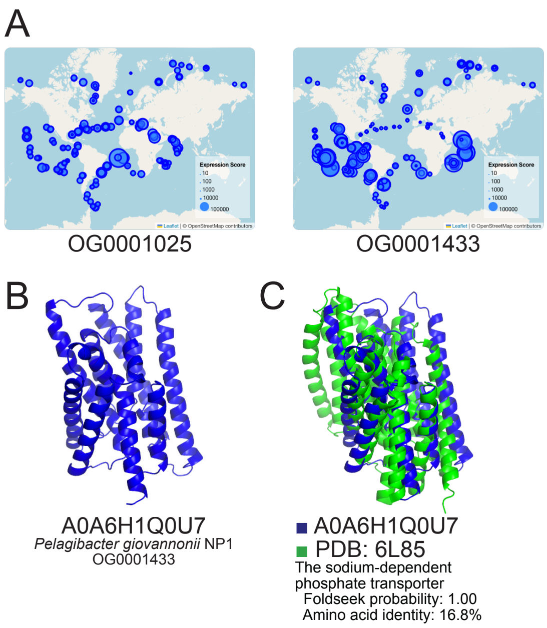


**Supplementary Figure S4. Environmental distribution and structural similarity of the alternative phosphate transporter-like OG0001433.**
(A) Tara Oceans Expression Score maps for the canonical phosphate transporter permease PstA (OG0001025, left) and the alternative phosphate-transporter-like OG0001433 (right). Symbol size represents the summed OG-level Expression Score. (B) AlphaFoldDB model for UniProt accession A0A6H1Q0U7 from *Pelagibacter giovannonii* NP1, a representative protein assigned to OG0001433. (C) Structural superposition of A0A6H1Q0U7 and the sodium-dependent phosphate transporter structure PDB 6L85 identified by Foldseek. The structural match had a Foldseek probability of 1.00 and an amino acid sequence identity of 16.8%, providing structure-based support for the proposed transporter function.

#### SUPPLEMENTARY TABLES

**Supplementary Table S1. Genome metadata for the 542 SAR11 genomes included in the Atlas.**

The table reports genome identifiers; clade, subclade, subgroup, family, genus, and species assignments; genome type; source references; sampling depth and coordinates; depth and latitude categories; Longhurst province assignments for marine samples; environmental and habitat descriptions; water-body information for non-marine samples; CheckM2 completeness and contamination estimates; assembly and coding statistics; nitrogen, carbon, and sulfur average residue side-chain composition values; and genomic GC content. The collection comprises 132 cultured strains, 399 SAGs, and 11 MAGs.

**Supplementary Table S2. CheckM2 quality assessment and genome-selection results for the SAR11 candidate pool.**

CheckM2 v1.0.2 quality estimates and assembly statistics are provided for 4,211 screened candidate *Pelagibacterales* genomes. Columns indicate whether each genome met the principal criteria of completeness >85% and contamination <10%, whether it was selected for the 542-genome Atlas, whether it was retained as a documented inclusion exception, and its final selection status. A total of 540 selected genomes met both quality thresholds; HIMB2304 and ETNP2013_S02_SV82_300m_MAG_01 were retained as documented exceptions.

**Supplementary Table S3. Functional annotation summary for the 4,577 SAR11 orthogroups.**

For each OG, the table reports the representative COG2024, KEGG Orthology, and Pfam assignments; COG gene names, functional letters, and categories; the number of assigned genes and genomes; genome prevalence; annotation support and coverage; top-assignment rates; and the number and coverage of Pfam domain types. Annotation summaries were calculated from all genes assigned to each OG.

**Supplementary Table S4. Feature-level comparison of the SAR11 Genome Atlas with representative genomic and environmental resources.**

Resources were compared with respect to functions relevant to the objectives of the SAR11 Genome Atlas. The comparison is not intended to provide a comprehensive evaluation of all functions offered by each resource. “OG” denotes ortholog group.

Resource features were evaluated using STRING 2025, GTDB Release 11-RS232, Ocean Gene Atlas v2.0, and Cyanorak v2.1, accessed on 30 July 2026.

**Supplementary Table S5. Correlations between SAR11 orthogroup expression and Tara Oceans environmental variables.**

The table contains 41,193 OG–environment combinations, representing all 4,577 OGs evaluated against nine environmental variables. Pearson correlations were calculated using raw and log1p-transformed OG-level Expression Scores, whereas Spearman rank correlations were calculated using untransformed Expression Scores. For each OG–environment combination, the table reports the number of samples with complete environmental data, the number with nonzero OG expression, Pearson's r and Spearman's ρ, the corresponding nominal P-values, and Benjamini–Hochberg-adjusted P-values, together with OG prevalence and functional annotations.

**Supplementary Table S6. Orthogroups with expression positively associated with absolute latitude.**

The table lists the 43 OGs selected in Case Study 1 using the criteria Spearman's rho >0.5, Benjamini-Hochberg-adjusted q <0.05, and detectable expression in at least 10 Tara Oceans metatranscriptomes. Correlation statistics, sample counts, OG prevalence, and COG2024, KEGG Orthology, and Pfam annotations are provided for each OG.

**Supplementary Table S7. Orthogroups and functional modules examined in the Case Study 2.**

The table defines the 22 OGs used to examine mutually exclusive phylogenetic distributions. OGs are grouped into Mn/Zn homeostasis, phosphate acquisition, iron acquisition, oxidative-stress response, and heme-biosynthesis modules. For each OG, the table provides its current and previous OG identifiers, proposed role, display label, genome prevalence, functional annotations, annotation support and coverage, and the evidence used for interpretation.

**Supplementary Table S8. Environmental correlations of the 22 orthogroups examined in Case Study 2.**

The table reports 198 Spearman rank correlations between the 22 selected OGs and nine environmental variables, extracted from the complete correlation results presented in Table S5. For each comparison, the table provides the number of samples, Spearman's

$\rho$, nominal P-value, and Benjamini–Hochberg-adjusted P-value. The figure dot field identifies associations marked with a dot in Figure 4E, corresponding to |Spearman's $\rho$| $\geq 0.3$ and a Benjamini–Hochberg-adjusted P-value $< 0.05$.
