## Supplemental Text for "SAR11 Genome Atlas: a genome and gene catalog for functional profiling of the most abundant bacterial clade in the ocean"

**Supplementary Text**

**Supplementary Methods**

**Exceptions to genome-quality filtering**

All 132 isolate genomes were retained, including HIMB2304, which had an estimated completeness of 81.32% and therefore fell slightly below the general completeness threshold (**Figure S1**) (Freel et al., 2025). MAGs serving as nomenclatural types for species name were evaluated individually. ETNP2013_S02_SV82_300m_MAG_01 (GCA_017640265.1), representing *Anoxypelagibacter denitrificans*, was retained because it is the only available genome for this species. Its completeness estimated by CheckM2 is 84.01%, slightly below the threshold applied here, whereas the original study reported 87.08% completeness using CheckM v1.0.13 (Ruiz-Perez et al., 2021). By contrast, the genome representing *Mesopelagibacter carboxydoxydans* (GCA_017640275.1), a member of subclade Ic (family *Mesopelagibacteraceae*), was excluded because its estimated completeness was substantially lower in both the original study (76.92% using CheckM v1.0.13) and the present analysis (77.45% using CheckM2) (Ruiz-Perez et al., 2021). Subclade Ic remained represented by ARS1 and MED605, which had estimated completeness values of 96.81% and 91.07%, respectively (**Tables S1 and S2**).

**Phylogeny inference based on previous classifications**

Average nucleotide identity (ANI) values calculated by Freel et al. (2025) were used to identify genomes that could be confidently assigned to the same species as the representative genomes. Genomes exhibiting ANI ≥ 95% to a species representative were assigned the corresponding species-level taxonomy and associated higher-level classifications. For genomes that were not in Freel et al. (2025), pairwise ANI was calculated among all 542 genomes using fastANI v1.34 (Jain et al., 2018). Reciprocal ANI estimates were averaged for all genome pairs. Species names and their associated genus, subclade, and family labels were transferred from previously named reference genomes only when ANI was ≥ 95%. AAI was also estimated using the aai_wf workflow in CompareM v0.1.2 (https://github.com/dparks1134/CompareM). All genomes assigned to species based on ANI showed an AAI ≥ 95% to their selected species representative. Although an ANI threshold of 95% may over-split ecologically coherent SAR11 populations (Freel et al., 2025; Zhao et al., 2025), we applied it deliberately as a stringent criterion for transferring existing nomenclature rather than defining ecological populations.

Family-, genus-, and subclade-level assignments were evaluated using the SAR11_165 marker-set phylogeny. The phylogeny was rooted using the 20 alphaproteobacterial genomes outside SAR11 as an outgroup and pruned to retain the 542 SAR11 genomes. Genomes classified directly in previous studies (Fernandes et al., 2025; Freel et al., 2025; Lanclos et al., 2023) or through species representatives information were used as primary seed genomes. Genomes classified using the ANI criterion were used as supplementary seeds.

To prevent taxonomic labels from being extrapolated beyond the phylogenetic ranges defined by classified genomes, crown boundaries were defined exclusively from seed genomes, and candidate genomes were evaluated only within these seed-defined crowns. For each taxon in each phylogeny, internal nodes descending from the overall seed most recent common ancestor (MRCA) were screened as candidate crown nodes. A node was considered eligible for topology-based classification when it contained at least two seeds assigned to the same taxon, contained no seeds assigned to a competing taxon at the same rank, was consistent with the assigned parent family, and had UFBoot support ≥95. Supported nested crowns were evaluated independently so that instability at the overall seed MRCA did not automatically exclude genomes located within smaller, well-supported seed-defined clades. When multiple eligible crowns contained the same candidate genome, the smallest eligible crown, defined as the one containing the fewest descendant genomes, was selected. Genomes assigned solely by tree topology were not reused as seeds. Because genus and subclade designations generally corresponded to the same phylogenetic lineages, topology-based genus and subclade assignments were evaluated jointly and assigned the same confidence category. Species names were not inferred from tree topology and were assigned only from direct literature evidence, species representative information, or the ANI criterion described above. Monophyly was evaluated using the is.monophyletic function from the R package ape v5.8-1 (Paradis & Schliep, 2019).

**Generation of database-distributed files**

A concatenated alignment of the bac120 single-copy gene set (bac120, 5,036 amino acid sites) was generated using GTDB-Tk v2.1.0 classify-wf (Chaumeil et al., 2022). The bac120 and SAR11_165 phylogenies were independently inferred from both alignments using FastTree v2.1.11 (Price et al., 2010) with the options -lg -gamma -boot 1000. A phylogenetic tree was also inferred with IQ-TREE2 v2.2.0.3 using the ModelFinder-selected Q.insect+F+I+R10 model for bac120 with 1,000 ultrafast bootstrap (UFBoot) and 1,000 Shimodaira-Hasegawa-like approximate likelihood ratio test (SH-aLRT) replicates (Kalyaanamoorthy et al., 2017; Hoang et al., 2018; Minh et al., 2020).

To facilitate user-driven assignment of genes to SAR11 orthologous groups (OGs), we generated and distributed a multiple sequence alignment and an HMM profile for each OG. Multiple sequence alignments and HMM profiles for each OG were generated by OrthoFinder using FAMSA v2.4.1 (Deorowicz et al., 2016) and hmmbuild in HMMER v3.4, respectively.

**Tara Oceans metatranscriptome metadata curation**

Environmental metadata associated with Tara Oceans samples were integrated for downstream visualization and correlation analyses. The complete metadata table, including all original fields, is retained unchanged on the Download page of the web interface to ensure transparency and reproducibility. However, several variables were excluded from downstream screening and the interactive comparison panel because the archived integrated values contained implausible zero values or were inconsistent with the corresponding station-level records in PANGAEA (https://doi.pangaea.de/10.1594/PANGAEA.836319). These variables were Carbon.total, CO3, HCO3, and Alkalinity.total. Fluorescence and Density were excluded because they were redundant with fluorescence colored dissolved organic matter (fCDOM) and Sigma-theta, respectively. Lower.size.fraction was excluded because it was constant across the included samples and therefore provided no informative variation. Negative calibrated Chlorophyll A values were retained in the supplementary table as archived but were treated as missing values in downstream plotting and correlation analyses. Pearson correlations were calculated using both raw and log1p-transformed OG-level expression scores, whereas Spearman rank correlations were calculated using untransformed expression scores. For each environmental variable and correlation method, P-values from 4,577 OG-level tests were adjusted separately using the Benjamini–Hochberg method to control the false discovery rate.

**Sampling location metadata curation**

Sampling location and reference information were manually curated from the NCBI BioSample database and the original publications to ensure consistency across genomes. Marine provinces were assigned from sampling latitude and longitude coordinates using the Longhurst Provinces version 4 (March 2010) polygon layer (https://www.marineregions.org/sources.php#longhurst). Sampling locations were assigned to the corresponding Longhurst provinces based on their latitude and longitude coordinates using a point-in-polygon procedure. Records outside the Longhurst polygons were manually reviewed and assigned where appropriate, whereas records without coordinates and freshwater genomes were left unassigned. Environmental categories were assigned based on the sampling depth and geographic coordinates of each genome. Sampling depths (`depth`) were classified into three categories: depth ≤ 100 m (0–100 m), 100 m < depth ≤ 200 m (100–200 m), and >200 m, and recorded as depth_cat. Sampling latitudes were classified into six categories: southern polar (<−66.5°), southern temperate (−66.5° to <−23.5°), southern tropical (−23.5° to <0°), northern tropical (0° to <23.5°), northern temperate (23.5° to <66.5°), and northern polar (≥66.5°), and recorded as latitude_cat. Categories were recorded as NA when the corresponding depth or latitude was unavailable.

**Web interface testing and quality assurance**

The web interface was tested on macOS 14.1 using Google Chrome v150.0.7871.187 and Apple Safari v17.1 (19616.2.9.11.7), and on iOS 18.6 using Google Chrome v150.0.7871.113 and Apple Safari. Core functions, including OG search, genome browsing, synteny visualization, phylogenetic tree exploration, environmental expression plotting, protein-structure visualization, and file download, were evaluated using the representative OGs OG0000236 and OG0001433 and 132 cultivated strain genomes. Internal links and downloadable files were manually checked for accessibility. Dataset-level consistency checks were performed to confirm agreement between information displayed in the web interface and the corresponding downloadable tables.

**Supplementary Results**

**Case study 1: High-latitude-associated expression pattern highlights potential cold-adaptation functions in SAR11**

OG0001344 encoded OpuAC/ProX, the substrate-binding component of a proline/glycine-betaine ABC transporter (ρ = 0.671; **Figure S3A**). Its elevated expression may reflect enhanced demand for compatible-solute uptake in polar environments, where cells must cope with osmotic fluctuations and cold-induced physiological constraints.

Two components of the translational machinery also showed positive associations with absolute latitude. OG0000463 encoded ribosomal protein S15/RpsO (ρ = 0.556; **Figure S3B**), whereas OG0000449 encoded the ribosome-binding factor RbfA (ρ = 0.537; **Figure S3C**). RbfA promotes maturation of the 30S ribosomal subunit and couples ribosome biogenesis to translation initiation, with genetic evidence supporting its importance during growth at low temperature (Sharma & Woodson, 2020). Similarly, deletion of *rpsO* causes cold sensitivity and defective 30S-subunit biogenesis in *Escherichia coli* (Bubunenko et al., 2006). The latitude-associated expression of these OGs is therefore consistent with an increased requirement for efficient ribosome assembly and translation under low-temperature conditions.

Furthermore, OG0001280, encoding an SrmB-family RNA helicase, was strongly correlated with absolute latitude (ρ = 0.604; **Figure S3D**). RNA helicases can resolve stable RNA secondary structures and facilitate ribosome assembly, processes that become particularly important following temperature downshift (Zhang & Gross, 2021). OG0000583, encoding the acyl carrier protein AcpP, was also preferentially expressed at high latitudes (ρ = 0.581; **Figure S3E**). Because acyl carrier proteins provide intermediates for fatty-acid biosynthesis, this association is consistent with remodeling of membrane lipid composition to maintain fluidity at low temperature.

**Case study 2: Phylogenetic profiling highlights contrasting functional strategies in SAR11**

**Other examples of mutually exclusive functional modules**

Among the many additional co-occurring and anti-occurring relationships identified by CORGIAS, we selected three representative systems to further illustrate the functional diversity captured by the network **(Figure 4E and Table S8)**. First, an FTR1-like Fe²⁺ permease (OG0002237) showed anti-occurrence with a Fe³⁺ ABC uptake module comprising a substrate-binding protein (OG0000801), permease (OG0000843), and ATPase (OG0000921). OG0002237 was preferentially expressed in colder, higher-latitude, and fCDOM-rich samples, whereas the Fe³⁺ ABC module showed weaker and less consistent environmental relationships.

Because the redox state and bioavailability of iron vary with oxygen concentration, salinity, organic ligands, particle dynamics, and other environmental factors, iron acquisition systems may be shaped by habitat-specific iron accessibility. This pattern was especially evident for the Fe(II) permease, which was enriched in the freshwater/brackish lineage *Fontibacteriaceae* in our dataset. This enrichment is consistent with Lanclos et al. (2023), who showed that SAR11 subclade IIIa is structured along salinity gradients and exhibits subgroup-specific iron acquisition traits. While Fe(III) ABC transporters are broadly conserved across IIIa, the high-affinity Fe(II) transporter efeU is restricted to IIIa.3 and freshwater LD12, and the Fe(II)/Zn efflux transporter fieF is likewise characteristic of IIIa.3 and LD12 (Lanclos et al., 2023). Together, these observations suggest that freshwater *Fontibacteriaceae* may be adapted not only to reduced salinity but also to iron regimes in which Fe(II) acquisition is comparatively advantageous. In contrast, lineages in more saline marine environments may rely more heavily on Fe(III) or ligand-bound iron acquisition strategies.

Second, the catalase-peroxidase KatG (OG0001150) showed anti-occurrence with a rubrerythrin-related module comprising OG0001232, OG0001234, and OG0001238. KatG expression was positively associated with temperature and negatively associated with depth, nitrate, and fCDOM. In contrast, the three rubrerythrin-module OGs were positively associated with oxygen and negatively associated with depth and nitrate. Catalase-peroxidases and rubrerythrins detoxify hydrogen peroxide through fundamentally different catalytic strategies, relying on a heme cofactor and a non-heme diiron center, respectively (Vlasits et al., 2010; Barreiro et al., 2023). Their anti-occurring distributions therefore suggest that the SAR11 clade may use alternative enzymatic systems for oxidative-stress defense.

Finally, HemG (OG0001786) and HemJ (OG0000854), two non-homologous protoporphyrinogen oxidases involved in heme biosynthesis, showed strong anti-occurrence (q = 3.05 × 10⁻²⁰). This relationship is consistent with lineage-specific replacement between non-orthologous enzymes performing the same biochemical step. Because protoporphyrinogen oxidation is linked to cellular redox metabolism, this replacement may also reflect differences in respiratory-chain context or electron-acceptor availability among SAR11 lineages. In *E. coli*, HemG functions through respiratory electron-transfer chains, using oxygen under aerobic conditions and alternative acceptors such as nitrate or fumarate under anaerobic conditions (Möbius et al., 2010). HemJ, by contrast, represents a distinct non-homologous type and is thought to have a different evolutionary origin (Kobayashi et al., 2014). Thus, this result suggests that some OG replacements may be shaped by lineage-specific redox physiology.

**References**

Barreiro, D. S., Oliveira, R. N. S., & Pauleta, S. R. (2023). Bacterial peroxidases – Multivalent enzymes that enable the use of hydrogen peroxide for microaerobic and anaerobic proliferation. *Coordination Chemistry Reviews*, *485*, 215114. https://doi.org/10.1016/j.ccr.2023.215114

Bubunenko, M., Korepanov, A., Court, D. L., Jagannathan, I., Dickinson, D., Chaudhuri, B. R., Garber, M. B., & Culver, G. M. (2006). 30S ribosomal subunits can be assembled in vivo without primary binding ribosomal protein S15. *RNA*, *12*(7), 1229–1239. https://doi.org/10.1261/rna.2262106

Chaumeil, P.-A., Mussig, A. J., Hugenholtz, P., & Parks, D. H. (2022). GTDB-Tk v2: Memory friendly classification with the genome taxonomy database. *Bioinformatics (Oxford, England)*, *38*(23), 5315–5316. https://doi.org/10.1093/bioinformatics/btac672

Deorowicz, S., Debudaj-Grabysz, A., & Gudyś, A. (2016). FAMSA: Fast and accurate multiple sequence alignment of huge protein families. *Scientific Reports*, *6*(1), 33964. https://doi.org/10.1038/srep33964

Fernandes, C., Haber, M., Layoun, P., Chiriac, M.-C., Bulzu, P.-A., Ghai, R., Kasalicky, V., Shabarova, T., Grossart, H.-P., Woodhouse, J., Piwosz, K., Alonso, C., Zanetti, J., Hamilton, D. P., Ngochera, M., Nakano, S., Okazaki, Y., & Salcher, M. M. (2025). Ecophysiology and global dispersal of the freshwater SAR11-IIIb genus Fontibacterium. *Nature Microbiology*, *10*(9), 2194–2206. https://doi.org/10.1038/s41564-025-02091-8

Freel, K. C., Tucker, S. J., Freel, E. B., Stingl, U., Giovannoni, S. J., Eren, A. M., & Rappé, M. S. (2025). New SAR11 isolate genomes and global marine metagenomes resolve ecologically relevant units within the Pelagibacterales. *Nature Communications*, *17*(1), 328. https://doi.org/10.1038/s41467-025-67043-6

Hoang, D. T., Chernomor, O., von Haeseler, A., Minh, B. Q., & Vinh, L. S. (2018). UFBoot2: Improving the Ultrafast Bootstrap Approximation. *Molecular Biology and Evolution*, *35*(2), 518–522. https://doi.org/10.1093/molbev/msx281

Jain, C., Rodriguez-R, L. M., Phillippy, A. M., Konstantinidis, K. T., & Aluru, S. (2018). High throughput ANI analysis of 90K prokaryotic genomes reveals clear species boundaries. *Nature Communications*, *9*(1), 5114. https://doi.org/10.1038/s41467-018-07641-9

Kalyaanamoorthy, S., Minh, B. Q., Wong, T. K. F., von Haeseler, A., & Jermiin, L. S. (2017). ModelFinder: Fast model selection for accurate phylogenetic estimates. *Nature Methods*, *14*(6), Article 6. https://doi.org/10.1038/nmeth.4285

Kobayashi, K., Masuda, T., Tajima, N., Wada, H., & Sato, N. (2014). Molecular Phylogeny and Intricate Evolutionary History of the Three Isofunctional Enzymes Involved in the Oxidation of Protoporphyrinogen IX. *Genome Biology and Evolution*, *6*(8), 2141–2155. https://doi.org/10.1093/gbe/evu170

Lanclos, V. C., Rasmussen, A. N., Kojima, C. Y., Cheng, C., Henson, M. W., Faircloth, B. C., Francis, C. A., & Thrash, J. C. (2023). Ecophysiology and genomics of the brackish water adapted SAR11 subclade IIIa. *The ISME Journal*, *17*(4), Article 4. https://doi.org/10.1038/s41396-023-01376-2

Minh, B. Q., Schmidt, H. A., Chernomor, O., Schrempf, D., Woodhams, M. D., von Haeseler, A., & Lanfear, R. (2020). IQ-TREE 2: New Models and Efficient Methods for Phylogenetic Inference in the Genomic Era. *Molecular Biology and Evolution*, *37*(5), 1530–1534. https://doi.org/10.1093/molbev/msaa015

Möbius, K., Arias-Cartin, R., Breckau, D., Hännig, A.-L., Riedmann, K., Biedendieck, R., Schröder, S., Becher, D., Magalon, A., Moser, J., Jahn, M., & Jahn, D. (2010). Heme biosynthesis is coupled to electron transport chains for energy generation. *Proceedings of the National Academy of Sciences of the United States of America*, *107*(23), 10436–10441. https://doi.org/10.1073/pnas.1000956107

Paradis, E., & Schliep, K. (2019). ape 5.0: An environment for modern phylogenetics and evolutionary analyses in R. *Bioinformatics*, *35*(3), 526–528. https://doi.org/10.1093/bioinformatics/bty633

Price, M. N., Dehal, P. S., & Arkin, A. P. (2010). FastTree 2 – Approximately Maximum-Likelihood Trees for Large Alignments. *PLOS ONE*, *5*(3), e9490. https://doi.org/10.1371/journal.pone.0009490

Ruiz-Perez, C. A., Bertagnolli, A. D., Tsementzi, D., Woyke, T., Stewart, F. J., & Konstantinidis, K. T. (2021). Description of Candidatus Mesopelagibacter carboxydoxydans and Candidatus Anoxipelagibacter denitrificans: Nitrate-reducing SAR11 genera that dominate mesopelagic and anoxic marine zones. *Systematic and Applied Microbiology*, *44*(2), 126185. https://doi.org/10.1016/j.syapm.2021.126185

Sharma, I. M., & Woodson, S. A. (2020). RbfA and IF3 couple ribosome biogenesis and translation initiation to increase stress tolerance. *Nucleic Acids Research*, *48*(1), 359–372. https://doi.org/10.1093/nar/gkz1065

Vlasits, J., Jakopitsch, C., Bernroitner, M., Zamocky, M., Furtmüller, P. G., & Obinger, C. (2010). Mechanisms of catalase activity of heme peroxidases. *Archives of Biochemistry and Biophysics, Heme Peroxidases*, *500*(1), 74–81. https://doi.org/10.1016/j.abb.2010.04.018

Zhang, Y., & Gross, C. A. (2021). Cold Shock Response in Bacteria. *Annual Review of Genetics*, *55*(Volume 55, 2021), 377–400. https://doi.org/10.1146/annurev-genet-071819-031654

Zhao, J., Pachiadaki, M., Conrad, R. E., Hatt, J. K., Bristow, L. A., Rodriguez-R, L. M., Rossello-Mora, R., Stewart, F. J., & Konstantinidis, K. T. (2025). Promiscuous and genome-wide recombination underlies the sequence-discrete species of the SAR11 lineage in the deep ocean. *The ISME Journal*, *19*(1), wraf072. https://doi.org/10.1093/ismejo/wraf072
